# The H2A.J histone variant contributes to Interferon-Stimulated Gene expression in senescence by its weak interaction with H1 and the derepression of repeated DNA sequences

**DOI:** 10.1101/2020.10.29.361204

**Authors:** Adèle Mangelinck, Clément Coudereau, Régis Courbeyrette, Khalid Ourarhni, Ali Hamiche, Christophe Redon, William M. Bonner, Erwin van Dijk, Céline Derbois, Robert Olaso, Jean-François Deleuze, François Fenaille, Claudia E. Rübe, Jean-Yves Thuret, Carl Mann

## Abstract

The histone variant H2A.J was previously shown to accumulate in senescent human fibroblasts with persistent DNA damage to promote inflammatory gene expression, but its mechanism of action was unknown. We show that H2A.J accumulation contributes to weakening the association of histone H1 to chromatin and increasing its turnover. Decreased H1 in senescence is correlated with increased expression of some repeated DNA sequences, increased expression of STAT/IRF transcription factors, and transcriptional activation of Interferon-Stimulated Genes (ISGs). The H2A.J-specific Val-11 moderates the transcriptional activity of H2A.J, and H2A.J-specific Ser-123 can be phosphorylated in response to DNA damage with potentiation of its transcriptional activity by the phospho-mimetic S123E mutation. Our work demonstrates the functional importance of H2A.J-specific residues and potential mechanisms for its function in promoting inflammatory gene expression in senescence.

## INTRODUCTION

Eucaryotic DNA is contained in nucleosomes composed of 2 copies each of 4 core histones (H2A, H2B, H3, H4), or their variants, plus a linker histone from the H1 family (1). Packaging of DNA into nucleosomes compacts the genome and restricts its access to enzymes involved in DNA transcription, replication, and repair. Expression of the canonical core histones is linked to the replication of DNA (2) and allows rapid reassembly of replicated DNA into chromatin. In contrast, a set of histone variants are typically expressed in a constitutive fashion that allows their continued expression in post-mitotic cells (3). Histone variants are thought to play specialized roles in gene transcription and DNA repair due to their specific protein sequences. For example, the H3.3 variant differs from canonical H3.1/H3.2 by only 4/5 amino acids that confer specificity with regards to recognition by histone chaperone complexes and by differential post-translation modification (4). The H2A.X variant has a short C-terminal extension containing an SQE motif that is phosphorylated by DNA damage response kinases in response to DNA damage. Phospho-H2A.X then recruits proteins involved in DNA repair and recombination (5).

The H1 linker histones join adjacent nucleosomes and contribute to chromatin compaction that can repress gene transcription (6). Indeed, depletion of histone H1 was shown to lead to derepression of repeated DNA sequences that subsequently triggered the expression of innate immunity pathways leading to expression of Interferon-Stimulated Genes (ISGs) (7). Association of H1 to chromatin involves its binding to both linker DNA and core histones including the C-terminal region of H2A (8, 9). The H2A.J histone variant encoded by the unique H2AFJ gene (or H2AJ gene in the new HUGO histone gene nomenclature) is expressed constitutively and differs by canonical H2A by Val-11 instead of Ala-11, and the last 7 amino acids containing an SQ motif in H2A.J (10, 11). We previously showed that H2A.J accumulates in senescent human fibroblasts induced into senescence by persistent DNA damage where it promotes inflammatory gene expression, an important component of the senescent-associated secretory phenotype (SASP) (11). H2A.J was found to be widely deposited in chromatin, and its mechanism of action was not defined. In this work, we show that H2A.J-specific residues are functionally important, and we show that H2A.J accumulation contributes to decreased levels of H1 in senescence, derepression of repeated DNA sequences, and activation of ISG expression.

## MATERIAL AND METHODS

### Cells and cell culture

WI-38hTERT/ pTRIPz-sh-No Target (Thermo RHS4743) and WI-38hTERT/pTRIPz-sh3-H2AFJ (Thermo V3THS_351902) human fetal pulmonary fibroblasts and their culture conditions were previously described (11). Cells were mycoplasma negative. Senescence was induced by etoposide treatment following a protocol that allowed rapid induction of inflammatory gene expression (12). Briefly, cells were grown in the presence of 1 μg/ml doxycycline for 3 days to induce the expression of sh-RNAs before senescence induction by incubation with 100 μM etoposide for 48 hours. Cells were then washed and incubated in fresh medium without etoposide for 5 days before harvesting cells for analysis. Doxycycline was maintained throughout the experiment to ensure continual expression of the sh-RNAs.

Stable WI-38hTERT cell lines that allow doxycycline-regulated expression of H2A.J, H2A.J mutants, and H2A-type1 were produced by infection with pTRIPz lentiviruses in which the appropriate H2A.J or H2A coding sequences were cloned as synthetic AgeI-MluI DNA fragments in place of the corresponding pTRIPz fragment. Transduced cells were selected for resistance to 1 μg/ml puromycin. The synthetic DNA also coded for a Flag-HA epitope that was added to the N-terminus of the histone proteins to allow their expression to be distinguished from endogenous sequences, and silent mutations rendered the encoded H2AFJ mRNAs resistant to the action of sh-H2AFJ RNAs. The sequence of the synthetic DNA fragments is shown in Supplementary Table 1. Lentiviruses were prepared from pTRIPz plasmids by co-transfection of 293T cells with packaging plasmids as described (13).

The effect of expressing H2A.J, H2A.J mutants, and H2A-type1 on the transcriptome of proliferating WI-38hTERT fibroblasts was performed by inducing the expression of each construct with 100 ng/ml doxycycline for 7 days before harvesting RNAs. Multiple genes encode canonical H2A species that are polymorphic at several amino acid residues (11). The H2A-type1 sequence chosen to represent canonical H2A in this study, in addition to Ala-11 and a canonical H2A C-terminus, differs from H2A.J by a threonine in position 17 instead of serine. However, other canonical H2A species also contain Ser-17 as for H2A.J. Thus, Ser-17 is not an H2A.J-specific residue.

T47D luminal breast epithelial cells for biochemical experiments were cultivated in RPMI-1640 + Gluta-Max medium containing 10% FBS + pen/strep.

### Mouse Embryonic Fibroblasts

A 7 bp deletion at the beginning of the H2AFJ gene was introduced by a TALEN-mediated DNA break in the C57Bl/6-N (Charles River) genome by Cyagen. This H2AFJΔ7 mutation created a frame shift with a premature stop codon at the beginning of the H2AFJ coding sequence (Supplementary Figure 1A). The founder heterozygous mutation was then back-crossed 6 times to C57BL/6-N mice provided by the Janvier Laboratory (France). Viable homozygous mutants were obtained after the second backcross. The homozygous H2A.J-ko mice are viable and fertile and their further phenotypic characterization will be described in a future publication. IF analysis of MEFs and mass spectrometric analysis of histones extracted from kidneys showed the absence of detectable H2A.J in the homozygous mutant (Supplementary Figures 1B and 1C). MEFs were produced from individual H2A.J-ko or isogenic WT C57BL/6-N embryos as described (14) with minor modifications as described below, and each embryo was sex genotyped by PCR (15). 3 different female WT and H2A.J-ko MEFs were used for the transcriptome analyses. MEFs were cultivated in DMEM + Gluta-Max (Gibco 31966) + 10% FBS + pen/strep in a 5% carbon dioxyde and 5% oxygen incubator. MEFs were passed twice before inducing senescence by treatment with 2.5 μM etoposide for 6 days (with media changed after 3 days), followed by 3 days incubation in fresh medium without etoposide (Supplementary Figures 1D and 1E). PolyA+ RNA-seq was performed as described below.

### Western blots

Whole cell extracts were prepared by flash freezing cell culture plates and scraping on ice in denaturing sample buffer containing protease and phosphatase inhibitors. Protein samples were slightly sonicated, heated to 95°C for 15 minutes, and electrophoresed in SDS-12% polyacrylamide gels and transferred to nitrocellulose membranes. Membranes were blocked in Intercept Blocking Buffer PBS (Eurobio 927-40003) and incubated overnight at 4°C with the first antibodies. The Li-Cor Odyssey CLx infrared imaging system was use to detect the second Dylight-680 or Dylight-800 labelled antibodies following the manufacturer’s protocols. The following antibodies were used for immunoblotting: anti-H2A.J (custom-made and characterized as described (11), 1:1,000), anti-H2A (Abcam, ab18255, 1:1,000), anti-H1.2 (Abcam, ab181977 and ab4086, 1:1,000 and 1:500), anti-flag M2 (Sigma, F1804, 1:1,000), anti-GAPDH (Abcam, ab9485, 1:5,000), anti-p16 (BD, 554079, 1:250), anti-p21 (Cell Signaling, 2947S, 1:1,000), anti-STAT1 (Invitrogen, AHO0832, 1:500), anti-phospho-STAT1 (Cell Signaling, 7649S, 1:250), anti-γH2AX (Millipore, 05-636, 1:1,000).

### Mass Spectrometric Identification of phospho-S123-H2A.J in irradiated mouse kidneys

A mouse was irradiated with 50 Gy ionizing irradiation, sacrificed 30 minutes post-IR, and organs were dissected and frozen. Histones were purified by acid extraction from kidneys and liver as previously described (11). To search for phosphorylation of H2A.J, acid-extracted histones were digested with Asp-N and Arg-C (Roche) and the resulting digests were analyzed by chromatography on an Ultimate 3000 nanoLC (Dionex, Sunnyvale, CA) coupled to a maXis UHR QTOF instrument (Bruker Daltonics, Bremen, Germany) equipped with a CaptiveSpray ionization source (Bruker Daltonics), essentially as described before (16). Peptides were first loaded onto a trapping column (C18 PepMap100, 300 μm i.d. × 5 mm, 5 μm) at 20 μL/min in 4% acetonitrile containing 0.1% formic acid. Then, the peptides were eluted towards the nano-column (C18 PepMap, 75 μm i.d. × 15 cm, 3 μm, 100 Å) at a flow rate of 300 nL/min, maintained at 35°C and using a 65-min linear gradient (from 4 to 35% acetonitrile containing 0.1% formic acid). The mass spectrometer source settings were as follows: capillary voltage, 1.5 kV; dry gas, 4.0 L/min; dry temperature, 150°C. The MS analysis was carried out in the positive ion mode with a resolution of ~50,000 at m/z ~1200, while MS/MS acquisitions were performed in a data-dependent acquisition mode using the seven most intense precursor ions. Precursor ions from the 400-2000 m/z window were fragmented under CID conditions for peptide identification. All data sets were re-calibrated using a lock mass (m/z 1221.9906) before data analysis. Resulting MS and MS/MS data were manually extracted and interpreted.

**Tandem affinity purification of Flag-HA-H2A.J and Flag-HA-H2A** were performed as previously described (17).

### Co-immunoprecipitation assays

Trypsinized cells were washed with PBS and quickly with Hypotonic Lysis Buffer (10 mM Tris-HCl (pH 7.5), 1.5 mM MgCl_2_, 10 mM KCl, 0.5 mM DTT, protease inhibitors). Cells were then incubated on ice for 10 minutes in Hypotonic Lysis Buffer and flash frozen in liquid nitrogen. Cell membranes were disrupted using a Dounce homogenizer with a type B pestle (25 strokes). Nuclei were pelleted by centrifugation at 1,500 g for 10 minutes at 4°C. Nuclear pellet was washed once with Hypotonic Lysis Buffer, then resuspended in Nuclear Extraction Buffer (25 mM Tris-HCl (pH 7.5), 500 mM NaCl, 1% Triton-X-100, 1% sodium deoxycholate, 1 mM DTT, protease inhibitors) and incubated on ice for 30 minutes. Chromatin and cell debris were pelleted by centrifugation at 21,000 × g for 15 minutes at 4°C and the supernatant was recovered as the nucleoplasmic fraction. This fraction was complemented with 25mM Tris-HCl (pH 7.5) to reduce NaCl concentration to 150 mM (600 μl final volume). Protein concentrations were determined using the Pierce 660 nm Protein Assay Kit (Thermo Scientific, #22662) and a NanoDrop 2000 spectrophotometer according to the manufacturer’s protocols. Fractions were treated with Benzonase Nuclease (Millipore, #E1014) for 30 minutes at room temperature. 10 μl of Protein A/G Magnetic Beads for IP (ThermoFisher Scientific, #10003D) were used for each IP. Beads were incubated for 1 hour at R.T. with 5 μg of antibody in 10 μl of IP Buffer (25 mM Tris-HCl (pH 7.5), 150 mM NaCl, 0.3% Triton-X-100, 0.3% sodium deoxycholate, 0.3 mM DTT, protease inhibitors). 10% BSA in IP Buffer were added to the beads for a 5% BSA final concentration. After a 2 hours incubation at 4°C, supernatants were removed and the antibody-coupled beads were incubated overnight at 4°C on a wheel with 200 μl / condition of nucleoplasmic fraction adjusted to equal protein concentrations (complemented with EGTA to 2 mM final concentration where indicated). After IP, beads were washed three times with 0.1% Tween-20 in PBS and finally resuspended in Western blot denaturing buffer. Samples were incubated for 15 minutes at 95°C for complete elution of proteins from the beads and analyzed by Western blot as described above. The following antibodies were used for IP: anti-IgG (Cell Signaling, 2729S), anti-H1.2 (Abcam, ab181977 and ab4086).

### Identification of H1.2-H2A complexes

Approximately 30 million T47D cells were were trypsinized and washed with PBS and then with Hypotonic Lysis Buffer (10 mM Tris-HCl (pH 7.5), 1.5 mM MgCl_2_, 10 mM KCl, 0.5 mM DTT, protease inhibitors). Cells were then incubated on ice for 10 minutes in Hypotonic Lysis Buffer and flash frozen in liquid nitrogen. Cell membranes were disrupted using a Dounce homogenizer with a type B pestle (25 strokes). Nuclei were pelleted by centrifugation at 23,000 × g for 5 minutes at 4°C. The nuclear pellet was washed once with Hypotonic Lysis Buffer then resuspended in Extraction Buffer (20 mM Tris-HCl (pH 7.5), 500 mM NaCl, 1.5 mM MgCl_2_, 2 mM EGTA, 0.5 mM DTT, 20% glycerol, protease inhibitors) and incubated for 1 hour at 4°C with agitation. Supernatant was recovered after centrifugation at 89,000 × g for 20 minutes at 4°C. This extract was kept on ice for 2 hours. Protein complexes were then separated by gel exclusion chromatography on a Superose 6 Increase 10/300 column with a 0.5 ml.min^−1^ flow rate of Extraction Buffer. 1/5 of 0.5 ml fractions were concentrated by TCA precipitation and analyzed by Western blot to identify H1.2 and H2A-containing fractions. Liquid chromatography coupled to tandem mass spectrometry of electrospray ionized tryptic peptides by the I2BC MS platform was used to identify histone chaperones within the gel filtration fractions containing H1.2 and H2A.

### Successive salt extractions of chromatin

Differential salt extraction of chromatin-associated proteins followed a previous protocol (18). Trypsinized cells were washed with PBS and quickly with Hypotonic Lysis Buffer (10 mM Tris-HCl (pH 7.5), 1.5 mM MgCl_2_, 10 mM KCl, 0.5 mM DTT, protease inhibitors). Cells were then incubated on ice for 10 minutes in Hypotonic Lysis Buffer and flash frozen in liquid nitrogen. Cell membranes were disrupted using a Dounce homogenizer with a type B pestle (25 strokes). Nuclei were pelleted by centrifugation at 1,500 × g for 10 minutes at 4°C. Nuclear pellets were washed once with Hypotonic Lysis Buffer then resuspended in Nuclear Extraction Buffer (25 mM Tris-HCl (pH 7.5), 500 mM NaCl, 1% Triton-X-100, 1% sodium deoxycholate, 1 mM DTT, protease inhibitors) and incubated on ice for 30 minutes. Chromatin was pelleted by centrifugation at 21,000 × g for 15 minutes at 4°C and the supernatant was recovered as the nucleoplasmic fraction. This fraction was complemented with 25mM Tris-HCl (pH 7.5) to reduce NaCl concentration to 150 mM and EGTA to 2 mM final concentration. The chromatin pellet was resuspended in MNase digestion Buffer (25 mM Tris-HCl (pH 7.5), 5 mM MgCl_2_, 1 mM CaCl_2_). Chromatin was incubated with 7.5 U.ml^−1^ MNase (Sigma, #N3755) for 15 minutes at 37°C with mixing at 800 rpm in an Eppendorf Thermomixer. The reaction was stopped by addition of EGTA to 2 mM final concentration and cooling on ice. Insoluble components were pelleted by centrifugation at 21,000 g for 15 minutes at 4°C and the supernatant was recovered as the MNase-digested fraction. This fraction was complemented with twice the fraction volume of IP Buffer (25 mM Tris-HCl (pH 7.5), 150 mM NaCl, 0.3% Triton-X-100, 0.3% sodium deoxycholate, 0.3 mM DTT, protease inhibitors) and EGTA to 2 mM final concentration. The pellet was resuspended in IP Buffer and incubate for 1 hour at 4°C on a wheel. Insoluble components were pelleted by centrifugation at 21,000 g for 15 minutes at 4°C. Supernatant was recovered as the 150 mM extract fraction. This fraction was complemented with EGTA to 2 mM final concentration. The pellet was resuspended in Max Extract Buffer (25 mM Tris-HCl (pH 7.5), 600 mM NaCl, 0.3% Triton-X-100, 0.3% sodium deoxycholate, 0.3 mM DTT, protease inhibitors) and incubated overnight at 4°C on a wheel. Insoluble components were pelleted by centrifugation at 21,000 g for 15 minutes at 4°C and the supernatant was recovered as the 600 mM extract fraction. This fraction was complemented with EGTA to 2 mM final concentration. All fractions were concentrated and desalted by TCA precipitation and analyzed by Western blot.

### H1 turnover following cyloheximide shutoff of protein synthesis

Cells were treated with 50μg/ml cycloheximide and flash frozen at indicated time points. Protein extraction and analysis were performed as described in the Western blot protocol.

### EdU incorporation to assess cell proliferation

5-ethynyl-2’-deoxyuridine (EdU) was added to cell culture medium to 25 nM final concentration for 24 hours. Cell fixation, permeabilization and EdU detection were performed with Click-iT EdU Alexa Fluor 488 Imaging Kit (ThermoFisher Scientific #C10337) according to the manufacturer’s protocols. Imaging was performed using a Leica DMIRE2 inverted motorized microscope.

### Senescence-associated β-galactosidase activity

Formaldehyde-fixed cells were stained with 1 mg/ml 5-bromo-4-chloro-3-indolyl-β-D-galactoside (X-gal) in 40 mM citric acid-sodium phosphate (pH 6.0), 5 mM potassium ferrocyanide, 5 mM potassium ferricyanide, 150 mM NaCl, 2 mM MgCl_2_ buffer for 40 hours at 37°C as described (19).

### Immunofluorescence

Formaldehyde-fixed cells were permeabilized with 0.2% Triton-X-100 in PBS, blocked with 5% BSA in 0.1% Tween-20 and PBS for 10 minutes and incubated with primary antibody in blocking buffer for 2 hours at room temperature. Cells were then incubated with an Alexa-488-labeled secondary antibody diluted to 1:500 in blocking buffer for 1 hour at room temperature. Nuclei were stained with 500 ng/ml DAPI for 10 minutes at room temperature. Imaging was performed using Metamorph software on a Leica DMIRE2 inverted motorized microscope. The following primary antibodies were used: anti-H2A.J (custom-made as previously described and characterized (11), 1:1,000), anti-p21 (Cell Signaling, 2947S, 1:500), anti-IL1α(R&D Systems, MAB200, 1:200), anti-IL6 (R&D Systems, AF-206, 1:200), anti-STAT1 (Invitrogen, AHO0832, 1:250).

### ChIP-seq

Cells were crosslinked with 1% formaldehyde for 10 minutes and then quenched with glycine at 200 mM final concentration. Cells were washed with cold PBS, scraped and pelleted by centrifugation at 1,500 × g for 10 minutes at 4°C. Cells were washed again twice with cold PBS and stored at −80°C in 50 μl of PBS. 150 μl of SDS Lysis Buffer (50 mM Tris-HCl (pH 8.0), 1% SDS, 1 mM EDTA, protease inhibitors) was added to crosslinked cells. Cells were incubated on ice for 10 minutes before transfer to 2 ml Eppendorf tubes and addition of 1,300 μl of MNase ChIP dilution buffer (20mM Tris-HCl (pH 8.0), 1% Triton-X-100, 1 mM EDTA, 150 mM NaCl, 3 mM CaCl_2_). Cells were pre-warmed to 37°C for 5 minutes before incubation with 1 U/ml of MNase (Sigma, #N3755) for 15 minutes at 37°C with mixing at 800 rpm in an Eppendorf Thermomixer. The reaction was stopped by addition of EGTA to 10 mM final concentration and cooling on ice. Cells were pelleted by centrifugation at 5,000 g for 5 minutes at 4°C. Cell pellets were resuspended in 300 μl of RIPA buffer (1X PBS, 1% NP-40, 0.5% sodium deoxycholate, 0.1% SDS, protease inhibitors) and transferred into Bioruptor-specific tubes. Sonication was performed with a Diagenode Bioruptor Pico sonication device using the following program: 10 cycles of 30 seconds on and 30 seconds off. Cell debris were pelleted by centrifugation at 15,000 × g for 5 minutes at 4°C. Chromatin fragmentation was checked by treating 5% aliquots of each step (bulk after MNase treatment, supernatant after MNase treatment and supernatant after sonication) with 0.1 mg/ml RNase A at 37°C for 1 hour and 0.2 mg/ml proteinase K at 65°C overnight, phenol/chloroform purification and then running in a 3% agarose gel in Sodium-Borate Buffer (10 mM sodium hydroxide, pH adjusted to 8.5 with boric acid) at 200 V for 30 minutes to check lengths of the DNA fragments. The protocol was optimized to yield a peak of fragmented chromatin around 100-300bp. DNA concentrations were determined using an Invitrogen Qubit 2.0 Fluorometer according to the manufacturer’s protocols. 100 μl of Dynabeads protein G (ThermoFisher Scientific, #10003D) were used for each IP. Beads were washed three times with 1 ml of PBS/BSA mix (5 mg/ml BSA in 1X PBS, protease inhibitors) before being resuspended in 300 μl of PBS/BSA mix and stored at 4°C overnight on a wheel. 5 μg of antibody was added to the beads and tubes were put back on the wheel at 4°C for 2 hours. The following antibodies were used: anti-H3K4me3 (Cell Signaling, 9751S), anti-RNA polymerase II (Abcam, ab817), anti-H3K27me3 (Millipore, 07-449) and anti-H3K9me3 (Abcam, ab8898). Antibody-coupled beads were washed three times with cold PBS/BSA mix before being resuspended in 100 μl of PBS/BSA mix. 300 μl of fragmented chromatin adjusted to equal DNA concentrations between conditions was added to the antibody-coupled beads and tubes were put back on the wheel at room temperature for 1 hour, then at 4°C for additional 2 hours. A 5% Input of fragmented chromatin was kept. Supernatant was discarded and beads were washed five times with 1 ml of cold LiCl Wash Buffer (100 mM Tris-HCl (pH 7.5), 500 mM LiCl, 1% NP-40, 1% sodium deoxycholate) for 3 minutes on a wheel at 4°C then once with 1 ml of cold TE Wash Buffer (10 mM Tris-HCl (pH 7.5), 0.1 mM Tris-EDTA) for 1 minute on a wheel at 4°C. Beads were resuspended in 200 μl of IP Elution Buffer (1% SDS, 0.1 M NaHCO_3_) and Input volumes were adjusted to 200 μl with IP Elution Buffer. Reverse crosslinking and ChIPed DNA recovery was performed by incubating the beads-and Input-containing tubes at 65°C for 1 hour with vortexing every 15 minutes. Beads-containing tubes were centrifuged at 21,000 g for 3 minutes at room temperature to recover the ChIPed DNA-containing supernatant. Each Input and ChIP sample was treated with 0.1 mg.ml^−1^ RNase A at 37°C for 1 hour and 0.2 mg.ml^−1^ proteinase K at 65°C overnight, then processed through phenol/chloroform purification. DNA concentrations were determined using the Qubit.

Illumina ChIP-seq libraries were then prepared with the NEBNext Ultra II DNA Library Preparation Kit (#E7645S) according to the manufacturer’s protocols. DNA concentration was checked using the Qubit after PCR amplification and after cleanup of PCR reaction. Size distribution was checked after cleanup of PCR reaction on an Agilent 2100 Bioanalyzer High Sensitivity DNA chip. Correct ligation of adaptors and sequencing primers was also checked by PCR and migration in a 3% agarose gel in Sodium-Borate Buffer at 200 V for 30 minutes. 43 bp paired-end DNA sequencing on an Illumina NextSeq 500 apparatus was performed by the I2BC high throughput DNA sequencing platform.

### ChIP-qPCR

Cells were crosslinked with formaldehyde 1% for 10 minutes and then quenched with glycine at 200 mM final concentration. Cells were washed with cold PBS, scraped and pelleted by centrifugation at 1,500 g for 10 minutes at 4°C. Cells were washed again twice with cold PBS and stored at −80°C in 50 μl of PBS. 300 μl of MNase-PBS Buffer (1X PBS, 1% NP-40, 3 mM CaCl_2_, protease inhibitors) was added to crosslinked cells and then transferred to Bioruptor-specific tubes. A first sonication was performed to liberate nuclei with a Diagenode Bioruptor Pico sonication device using the following program: 6 cycles of 10 seconds on and 30 seconds off. This program had to be repeated once for senescence conditions. Tubes were pre-warmed to 37°C for 5 minutes before incubation with 10 U/ml MNase (Sigma, #N3755) for 15 minutes at 37°C under spinning at 800 rpm in an Eppendorf Thermomixer comfort. The reaction was stopped by addition of EGTA to 10 mM final concentration and cooling on ice. A second sonication was performed to liberate the fragmented chromatin with a Diagenode Bioruptor Pico sonication device using the following program: 6 cycles of [30 seconds ON – 30 seconds OFF]. MNase-PBS Buffer complemented with SDS and sodium deoxycholate, respectively to 0.1% and 0.5% final concentration. Cell debris were pelleted by centrifugation at 21,000 g for 10 minutes at 4°C. Each step of the fragmentation (initial state, liberation of nuclei and liberation of chromatin) was checked under a microscope.

Chromatin fragmentation was checked as described in ChIP-seq protocol and ChIP was performed as described in ChIP-seq protocol. The antibody used for ChIP was anti-H1.2 (Abcam, ab4086). Quantitative PCR was performed as described in RT-qPCR protocol. Values were normalized to Input DNA levels. DNA primers for the qPCR are shown in Supplementary Table 2.

### Total RNA-seq and small RNA-seq

RNA was purified using the mirVana miRNA Isolation Kit (ThermoFisher Scientific, #AM1560) according to the manufacturer’s protocols to yield a fraction containing total RNAs and a second fraction enriched in small RNAs. The total RNA fraction was used with the Illumina Ribo-Zero Scriptseq library kit to deplete cytosolic rRNA and prepare DNA librairies. The small RNA fraction was used with the NEBNext small RNA library prep kit (New England Biolabs, #E7300) to produce a small RNA library. Libraries were subjected to size selection on a 6% native polyacrylamide gel to enrich for molecules with inserts within the miRNA size range (18-25 nt). Both librairies were sequenced using 43 bp paired-end reads. The library preparation and DNA sequencing were performed by the I2BC high-throughput DNA sequencing platform.

### polyA+ RNA-seq

Cells were lysed and RNAs were purified using the Machery/Nagel Nucleospin RNA+ kit. The Illumina TruSeq Stranded mRNA kit was used to select polyA+ RNAs and to prepare DNA libraries for sequencing using 43 bp paired-end reads.

### RT-qPCR

RNA was extracted as indicated in RNA-seq and Illumina Bead ChIP Array protocols. cDNA was prepared from 500ng of RNA using random hexamer primers. Quantitative PCR was performed with a Luminaris Color HiGreen Fluorescein qPCR Master Mix (Thermo Scientific #K0384) and a CFX96 Touch Real-Time PCR Detection System (Bio-Rad) following the manufacturer’s protocols. Values were normalized to GAPDH RNA levels. DNA primers for the qPCR are shown in Supplementary Table 2.

### Illumina Bead ChIP Array transcriptome analyses

The effect of expressing H2A.J, H2A.J mutants, and H2A-type1 on the transcriptome of proliferating WI-38hTERT fibroblasts was assessed by microarray analyses using Illumina HumanHT-12v4 Bead Chip Arrays as previously described (11).

### DNA Sequence availability

RNA-seq, ChIP-seq, and microarray data were deposited in the EBI Array Express database under the following accession numbers: MEF polyA+ RNA-seq (E-MTAB-9704), Illumina Bead Chip microarrays (E-MTAB-9706), RNA-seq of small RNAs (E-MTAB-9710), RNA-seq of rRNA-depleted total RNAs (E-MTAB-9714), ChIP-seq of RNA polymerase II (E-MTAB-9717), ChIP-seq of H3-K4me3 (E-MTAB-9718).

### Bioinformatic analyses

. R and bash scripts are posted on Zenodo. Briefly, fastq files were trimmed with Cutadapt (20), quality controlled with FastQC (21), mapped to the hg19 human genome with bwa (22), or bowtie, or bowtie2 (23). For ChIP-seq experiments, peaks were identified with macs2 (24), annotated with ChIPseeker (25), and pathway enrichment analysis was performed with ReactomePA (26). For some RNA-seq analyses, alignment was performed to the transcriptome instead of the genome with salmon (27). Read counts were then aggregated to the gene level with tximeta (28), and differential gene expression was analysed with DESeq2 (29), edgeR (30), and limma-voom (31, 32). Gene set enrichment analysis was performed with camera (33) and with the Broad Institute server (34). RepEnrich2 (https://github.com/nerettilab/RepEnrich2) was used to assign bam files generated by bowtie2 to repeated DNA elements, and differential expression analysis was then performed as above for unique sequences.

The bash and R codes used for the ChIP-seq, Ribo-Zero RNA-seq, and small-RNA-seq analyses are availabe at Zenodo (10.5281/zenodo.4147010). Code and salmon quant files for the MEF transcriptome analysis are available at Zenodo (10.5281/zenodo.4140080). Illumina Bead Chip microarrray analysis code and idat files are available at Zenodo (doi:10.5281/zenodo.4139402).

## RESULTS

### H2A interacts more strongly with H1 compared to H2A.J

We immuno-purified Flag-HA-H2A.J from the nucleosoluble fraction of HeLa S3 cells to compare its interacting partners with those of similarly purified canonical H2A (Figure 1A and Supplementary Table 3). Most abundant proteins co-purifying with H2A.J also co-purified with H2A including Importin9 and the histone chaperones NAP1L and FACT (composed of subunits SPT16 and SSRP1) (Supplementary Table 3, highlighted in yellow). It is thus likely that H2A.J is imported into the nucleus and deposited in chromatin similarly to H2A. The only difference in identified bands between the H2A and H2A.J purifications was the presence of histone H1 in association with nucleosoluble H2A, but not H2A.J. We thus sought to verify this differential interaction using targeted co-immunoprecipitation experiments. We prepared WI-38hTERT human fibroblast cell lines expressing either Flag-HA-H2A.J or Flag-HA-H2A and immunoprecipitated histone H1 from the nucleosoluble fraction and evaluated the amount of co-precipitated H2A.J or H2A by immunoblotting. We observed significantly lower levels of H2A.J associated with H1 compared to H2A in 4 independent experiments (Figure 1B). Remarkably, we noticed that the interaction of H1 with H2A, and to a lesser extent with H2A.J, was increased in the presence of the calcium chelator EGTA. This result suggests that calcium, previously shown to affect the secondary structure of histone H1 (35), weakens the interaction of H1 with H2A and H2A.J. H2A is typically found in a heterodimeric complex with H2B (36, 37). Superose 6 gel filtration of T47D cell nucleosoluble extracts showed that H2A and H1 were found in high molecular weight fractions that also contained the histone chaperones NAP1L, NASP, and SSRP1 (Supplementary Figure 2 and Supplementary Table 4). It thus possible that H1 interacts with H2A-H2B in the nucleosoluble compartment of cells in association with histone chaperones.

**Figure 1.**
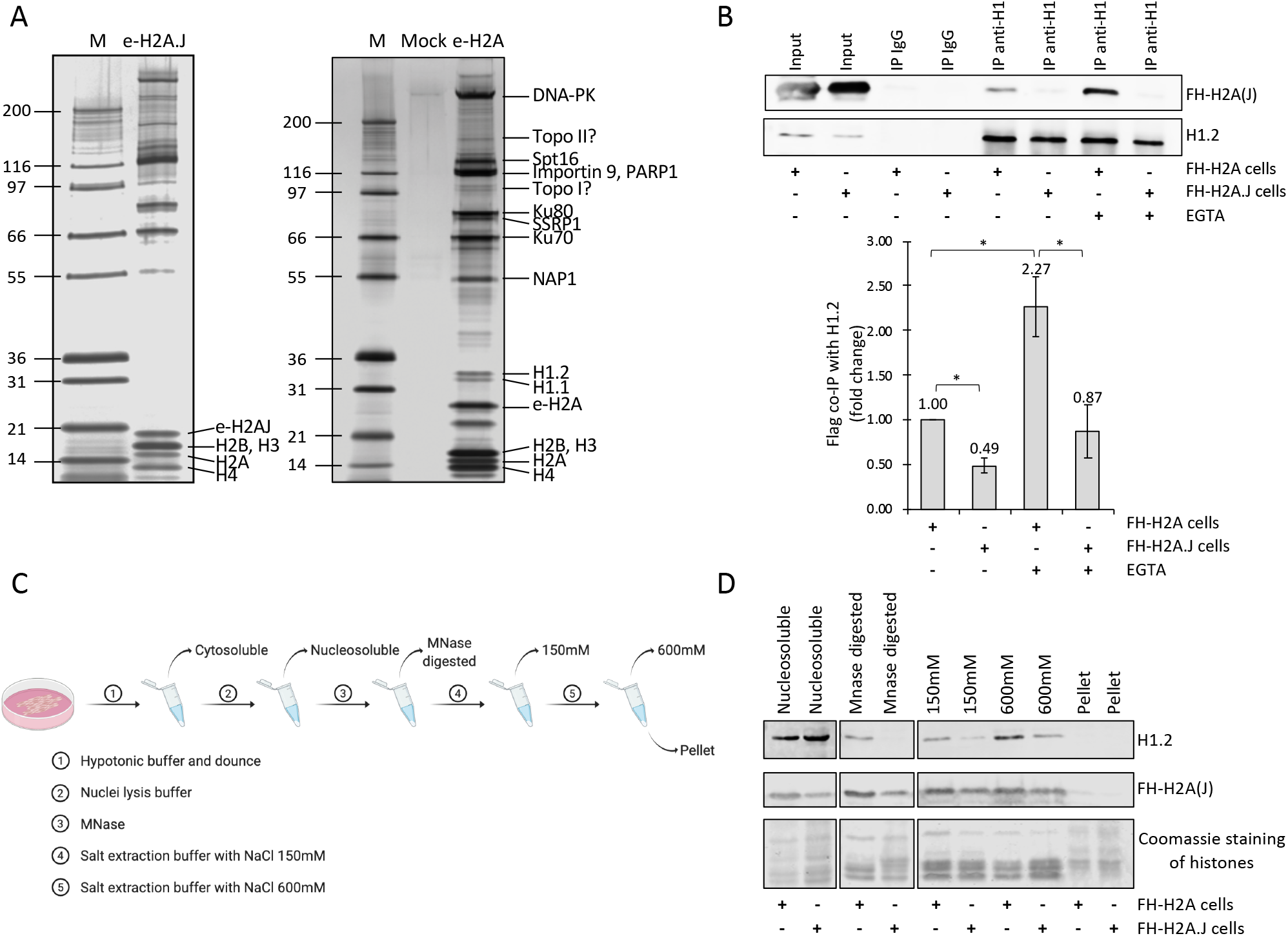
H1 interacts less strongly with H2A.J compared to H2A. **(A)** Tandem affinity purification of (Flag-HA)-epitope tagged H2A.J (e-H2A.J) and e-H2A from the nucleosoluble fraction of HeLa S3 cells. Shown is the Coomassie Blue stain of proteins eluted from the affinity beads. Bands identified by mass spectrometry are labeled. Lane M contains a molecular weight marker. The Mock lane represents a purification from HeLa cells that do not express an epitope-tagged protein. Note that the major difference between the e-H2A.J and e-H2A purifications is the apparent absence of H1 in the e-H2A.J purification. e-H2A migrates more slowly than e-H2A.J because it contained 2 copies of the N-terminal Flag-HA epitope. **(B)** Upper panel: Western blots of the co-IP of FH-H2A.J or FH-H2A with histone H1.2 from nucleosoluble extracts of WI-38hTERT fibroblasts expressing FH-H2A.J or FH-H2A, in the absence or the presence of EGTA. Input shows the amounts of FH-H2A, FH-H2A.J, and H1.2 in the nucleoplasmic fractions used to immunoprecipitate H1.2. Lower panel: quantification of the results of 4 independent co-IP experiments showing that H2A co-immunoprecipitates to a greater extent with H1.2 than does H2A.J, and both interactions are reinforced when the IP is done in the presence of EGTA. Asterisk, p < .05 for 2-sided t-test. **(C)** Schematic for the experiment involving differential salt extraction of chromatin to assay the quantity of H1.2 bound to chromatin. **(D)** Western blot showing increased amounts of H1.2 associated with the chromatin of WI-38hTERT fibroblasts overexpressing FH-H2A compared to cells overexpressing FH-H2A.J.

### H2A.J Overexpression Decreases H1 Levels

The lower apparent affinity of H2A.J for H1 suggests that increasing H2A.J deposition in chromatin might lead to a decreased chromatin association of H1. We explored this possibility by isolating nuclei from proliferating WI-38 fibroblasts over-expressing Flag-HA-H2A or Flag-HA-H2A.J and extracting chromatin-associated proteins by micrococcal nuclease (MNase) digestion and successive extraction with buffers containing increasing salt concentrations. MNase digestion in low-salt conditions preferentially liberates euchromatic nucleosomes whereas higher salt extractions are required to liberate heterochromatic nucleosomes (18). Histone H1 was associated to a greater extent with the chromatin of cells over-expressing H2A compared to H2A.J suggesting that its chromatin affinity and/or abundance is decreased by H2A.J overexpression (Figure 1C and 1D). Since H2A.J interacts less well with H1 compared to H2A, we considered that increased H2A.J levels in senescent fibroblasts might contribute to increased soluble H1 and a potentially increased turnover of H1. We first compared the effect of overexpressing Flag-HA-H2A.J versus Flag-HA-H2A in proliferating fibroblasts on total H1 levels. In three independent experiments, overexpression of H2A and H2A.J led to decreased levels of H1 in total cell extracts, but H2A.J overexpression led to greater decreases in H1 levels (Figure 2A). Cycloheximide protein synthesis shutoff experiments indicated that the half-life of H1 was shorter in fibroblasts over-expressing H2A.J compared to H2A (Figure 2B), which could explain the decreased H1 levels in cells over-expressing H2A.J.

**Figure 2.**
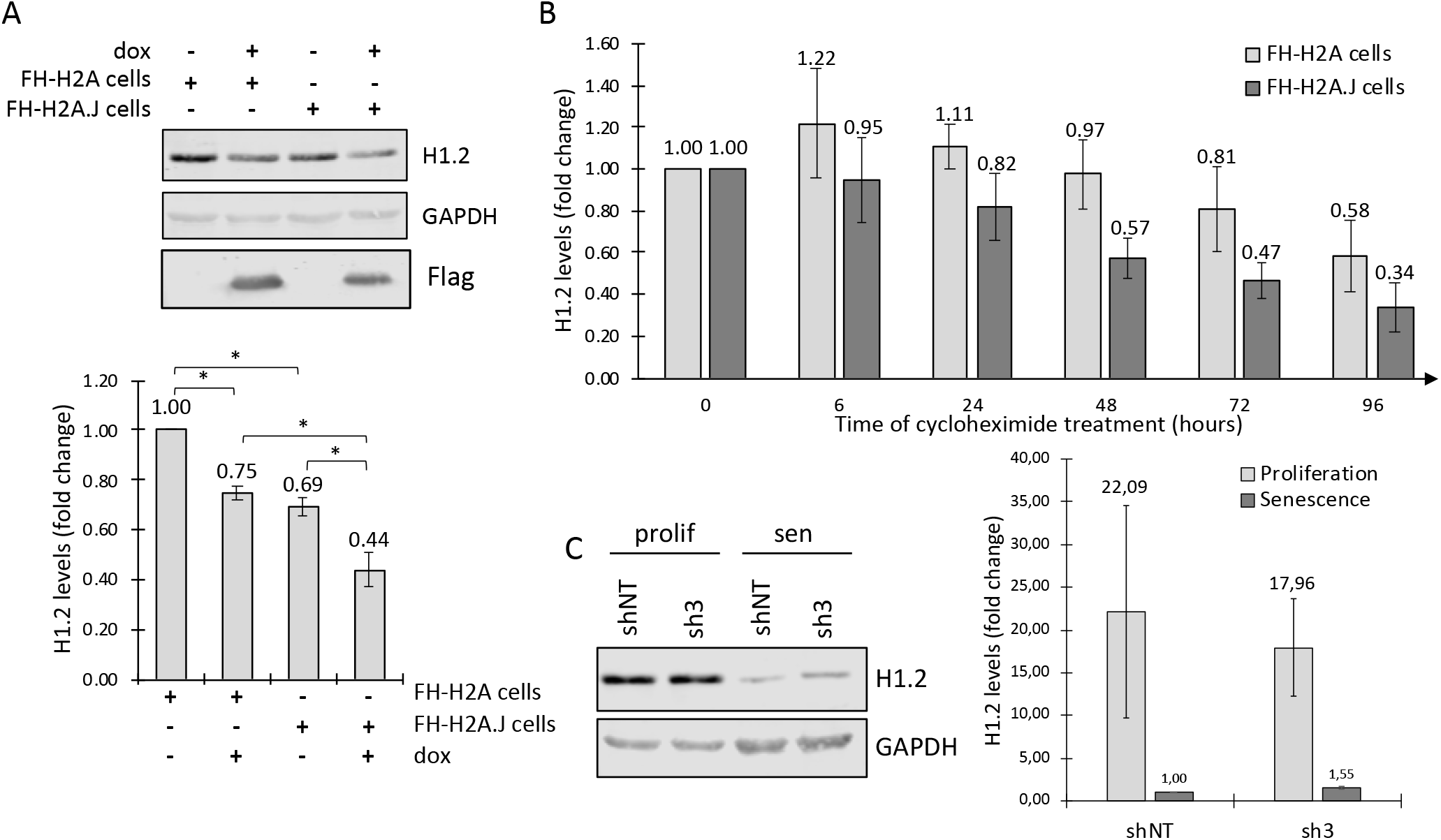
H2A.J overexpression decreases H1.2 levels and half-life. (**A**) Upper panel: representative H1.2 Western blot (n=3) showing that overexpression of H2A.J in WI-38hTERT fibroblasts decreases the total amount of H1.2 to a greater extent than overexpression of H2A. GAPDH was used as the normalization control. Lower panel: quantification of H1.2/GAPDH signal for 3 Western blots. Asterisk: p < .05 for 2-sided t-test. (**B**) Time-course of H1.2 levels in WI-38hTERT fibroblasts overexpressing H2A or H2A.J after addition of cycloheximide to block protein synthesis. Shown is the quantification of the H1.2/GAPDH signal ratio for 3 independent experiments. (**C**) Left panel: Western blot showing H1.2 levels in WI-38hTERT/sh-NoTarget and WI-38hTERT/sh3-H2AFJ fibroblasts in proliferation and in senescence induced by etoposide treatment. Right panel: Quantification for 2 independent experiments of H1.2 levels normalized to GAPDH and to the levels in shNT cells in senescence.

We previously showed that H2A.J levels are low in proliferating WI-38 fibroblasts and increased in senescent WI-38 fibroblasts (11). It was also shown that H1 levels decrease in senescent WI-38 fibroblasts (38). Consistent with a role for H2A.J in decreasing H1 levels, we found H1 levels to be slightly higher in senescent WI-38 fibroblasts depleted for H2A.J compared to control sh-NT cells (Figure 2C). The relatively modest difference in H1 levels suggests that other factors contribute to H1 decreases in senescence, such as transcriptional repression of H1 genes (see below), and increased calcium in senescence.

### H2A.J Expression Contributes to Interferon-Stimulated Gene Expression in Senescence

Our biochemical experiments above suggest that increased expression of H2A.J in senescent fibroblasts could contribute to the decrease of histone H1 in senescence. Previous work from the Jordan lab has shown that H1 depletion can lead to the derepression of Interferon-Stimulated Genes (ISGs) (7). Our previous work showed that H2A.J depletion inhibited inflammatory gene expression in senescent fibroblasts, and ectopic expression of H2A.J in proliferating fibroblasts stimulated inflammatory gene expression, and in particular, ISGs (11). Thus, decreased H1 levels in senescent fibroblasts, dependent in part on increased levels of H2A.J in senescence, could contribute to the transcription of ISGs as part of the inflammatory gene expression program in senescence. To test this possibility further, we performed genome-wide analyses of the activating histone H3-K4me3 activation mark and RNA pol II occupancy in WI-38/sh-NT and WI-38/sh-H2AFJ in proliferating and senescent fibroblasts (Figure 3). We induced senescence with etoposide using a protocol (12) that allowed rapid induction of inflammatory genes (Supplementary Figure 3). Macs2 (24) was used to identify ChIP-seq peaks that distinguished the sh-NT and sh-H2AFJ fibroblasts in etoposide-induced senescence. Peaks with differential intensity were mapped to proximal genes with ChIPseeker (25), and the generated gene set was then analyzed by Reactome pathway enrichment analysis (26). Interferon signaling pathways and Interferon-Stimulated Genes were the highest scoring pathways (Figure 3A, B, and C). Notably, MIR3142HG encoding mIR-146A, was the gene whose H3-K4me3 peak was the most affected by H2A.J knock-down (Figure 3D). RNA-seq of small RNAs showed that mIR-146A was highly induced in senescent fibroblasts (Figure 3E) as has been previously reported (39). Consistent with the decreased levels of the activating H3-K4me3, miR-146A levels were greatly reduced in H2A.J knock-down cells relative to the NT controls (Figure 3F and Supplementary Table 5). Interestingly, most previous data on miR-146A indicate that it is a negative feedback regulator of the Interferon pathway (39, 40), thereby suggesting a defect in this pathway in cells depleted for H2A.J.

**Figure 3.**
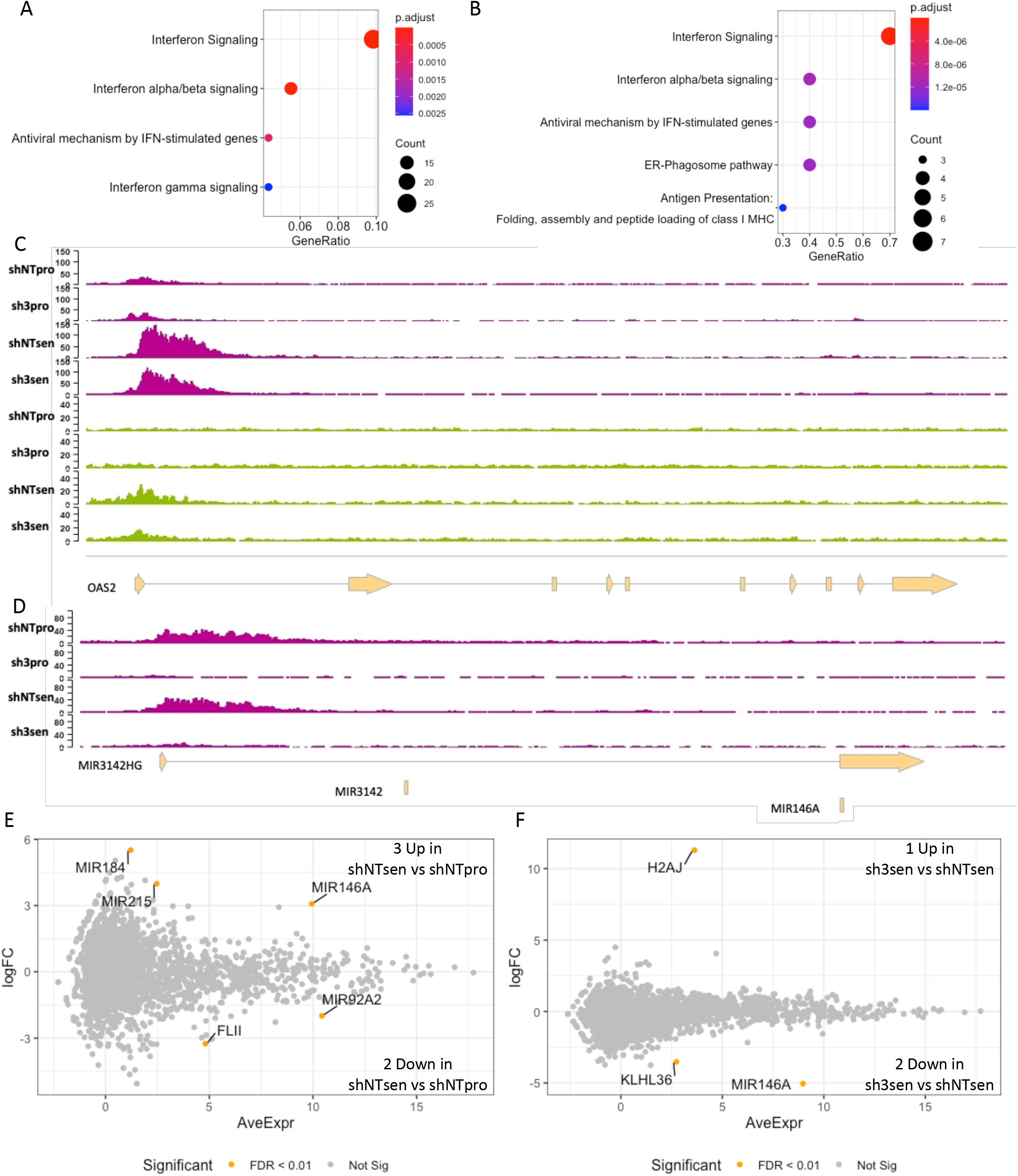
H3-K4me3 and RNA poymerase II ChIP-seq experiments show that interferon pathways are differentially affected by H2A.J knock-down in senescence, and are associated with defective induction of miR-146A, a negative feedback regulator of the interferon pathway. (**A,B**) Pathway enrichment analysis on differential H3-K4me3 peaks (A) and RNA polymerase II peaks (**B**). For both analyses, peaks that were both increased in senescent versus proliferating sh-NT cells, and in senescent sh-NT cells versus senescent sh-H2AFJ, were used for the Reactome pathway analysis. (**C**) Genome browser displays of RNA polymerase II ChIP-seq reads (magenta) and H3-K4me3 reads (citrus green) mapping to the OAS2 gene that is know to be activated by interferon. (**D**) Genome Browser displays of H3-K4me3 ChIP-seq reads mapping to the MIR3142HG encoding mIR-146A. (**E**) Log2FC-Average Expression plots of differentially expressed small RNAs for shNT cells in senescence versus proliferation. (**F**) Log2FC-Average Expression plots of differentially expressed small RNAs for shNT cells in senescence versus sh-H2AFJ cells in senescence. The H2AJ sequences up-regulated in sh3-H2AFJ cells are small RNA degradation products derived from the H2AFJ gene.

H1 depletion was associated with transcriptional derepression of repeated DNA sequences and transcriptional activation of ISGs (7). We previously reported transcriptome analyses of WI-38/hTERT fibroblasts expressing sh-NoTarget and two different H2AFJ-shRNAs using Illumina Bead Cheap arrays that showed a defect in inflammatory gene expression in H2A.J-deleted fibroblasts in senescence (11). We now describe Ribo-Zero RNA-seq to more globally assess effects of H2A.J depletion on RNA levels in senescence. Principal component analysis showed good separation of senescent versus proliferating and sh-NT versus sh-H2AF transcriptomes (Figure 4A). Comparison of sh-NT cells induced into senescence with etoposide relative to proliferating cells showed the anticipated repression of E2F-activated cell cycle genes and loss of canonical histone gene expression, including histone H1 genes, accompanied by increased expression of NFkB and Interferon-regulated inflammatory genes (Figure 4B, 4C and Supplementary Table 6). Comparison of sh-H2AFJ cells in senescence relative to sh-NT control cells in senescence also confirmed the previous microarray analysis indicating a defect in inflammatory gene expression in H2A.J knock-down cells (Figure 4D-G). GSEA indicated that down-regulation of Interferon Response Genes was the most significantly affected gene set (Figure 4E), which is consistent with the CHIP-seq results for H3-K4me3 and RNA polII (Figure 3). We next compared levels of RNAs transcribed from repeated DNA sequences. Consistent with previous studies (41, 42), most differentially expressed repeat RNAs were up-regulated in senescent relative to proliferating cells with the greatest increase assigned to HSATII pericentromeric satellite sequences (Figure 5A). Comparing H2A.J-ko cells with sh-NT cells in senescence, we found lower levels of RNAs transcribed from 7 repeated DNA elements and increased transcription from 4 elements (Figure 5B). Three repeated DNA elements, MER53, LTR26, and LTR23, were induced during the senescence of sh-NT control cells and underexpressed in the H2A.J knock-down cells in senescence relative to sh-NT cells. We identified DNA primers that allowed specific PCR amplification of LTR23 and LTR26 sequences from genomic DNA, but we were unsuccessful in finding primers that allowed amplification of MER53 sequences. We used the LTR23 and LTR26 primers in ChIP-qPCR experiments to quantify the abundance of H1.2 bound to those sequences in sh-NT and sh-H2AFJ cells in proliferation and in senescence. We found higher levels of H1.2 bound to LTR23 and LTR26 sequences in the H2A.J-knock-down cells compared to the sh-NT controls in both proliferation and senescence (Figure 5C). In the H2A.J knock-down cells relative to sh-NT, we also found higher levels of H1 bound to the promoter regions of the inflammatory genes IFI6, OAS2, CXCL8, and to the TYR gene that is not expressed in fibroblasts. For most of these sequences, there was less H1 bound in senescent cells compared to proliferating cells. Overall, these ChIP-qPCR results were correlated with global levels of H1 that were higher in sh-H2AFJ cells compared to sh-NT cells, and higher in proliferating compared to senescent cells (Figure 2C).

**Figure 4.**
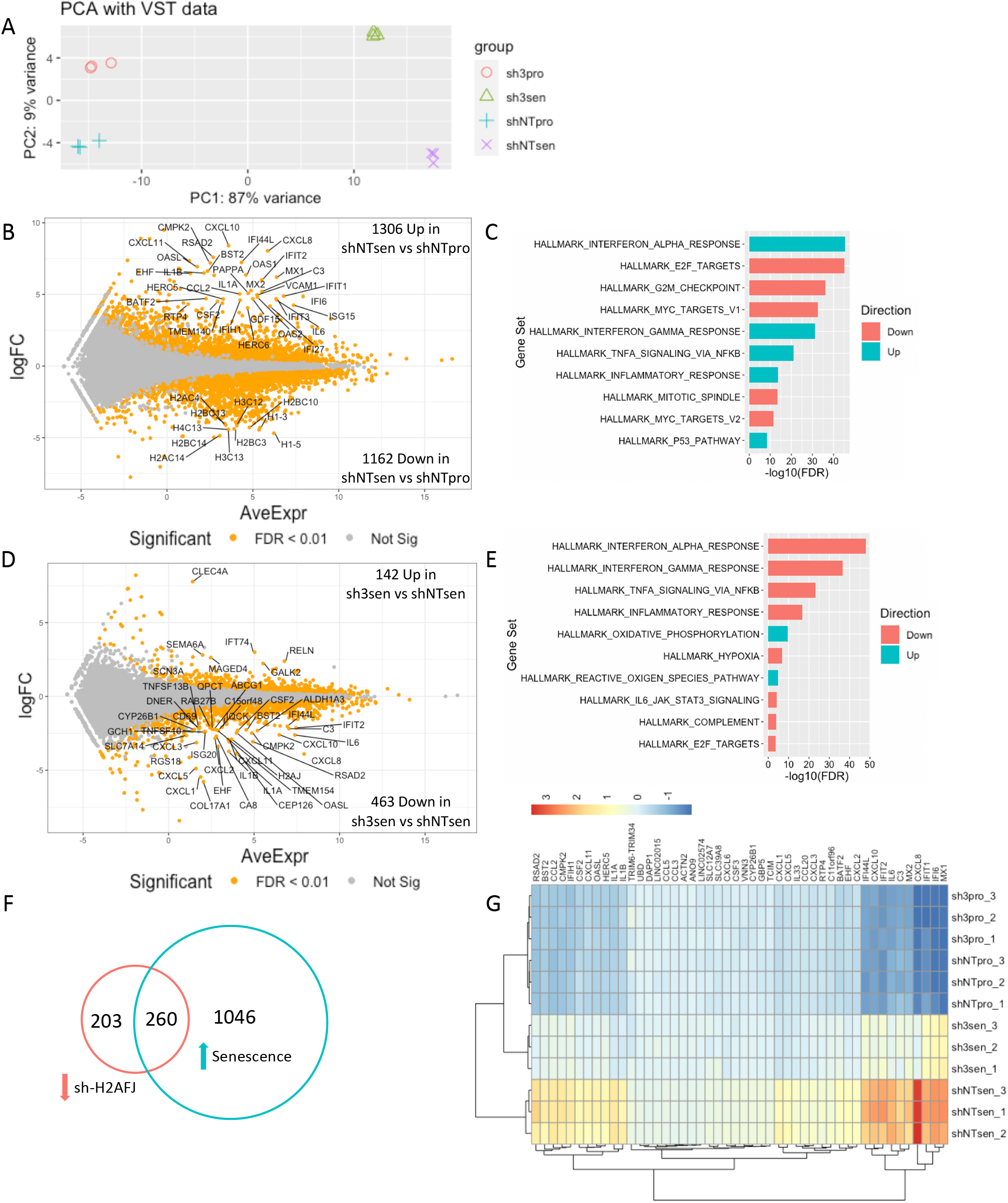
Ribo-Zero RNA-seq shows a defect in Interferon-Stimulate Gene expression in fibroblasts depleted for H2A.J in senescence. (**A**) PCA plot of the transcriptomes for 3 replicates each of WI-38hTERT/sh-NT and WI-38hTERT/sh3-H2AFJ cells showing good separation of senescent versus proliferating (PC1) and sh-NT versus sh3-H2AF transcriptomes (PC2). (**B**) Log2FC-Average Expression plots and (**C**) Hallmark GSEA of differentially expressed mRNAs for sh-NT cells in senescence versus proliferation. The GSEA shows highly significant Up-regulated enrichment of the senescent transcriptomes with Interferon and Inflammatory Response gene sets, and highly significant Down-regulated enrichment with E2F Target, G2M Checkpoint, and Myc Target gene sets as expected for senescent cells. (**D**) Log2FC-Average Expression plots of differentially expressed mRNAs for shNT cells in senescence versus sh-H2AFJ cells in senescence. (**E**) Log2FC-Average Expression plots and (**F**) Hallmark GSEA of differentially expressed mRNAs for senescent sh-H2AFJ cells versus senescent sh-NT cells. The GSEA shows highly significant enrichment with Down regulation of Interferon and Inflammatory Response gene sets in the sh3-H2AFJ cells in senescence. (**F**) Venn diagram summarizing the differential expression analysis indicating 1046 up-regulated genes in senescent versus proliferative sh-NT cells and 463 genes down-regulated in senescent sh-H2AFJ cells versus senescent sh-NT cells of which 260 represent genes that are also up-regulated in senescence in sh-NT cells. (**G**) Gene expression heat map for genes that are up-regulated (log2FC > 4) in senescent versus proliferative sh-NT cells and up-regulated in senescent sh-NT cells versus sh-H2AFJ cells (log2FC > 1.5) that is enriched for inflammatory and interferon-stimulated genes.

**Figure 5.**
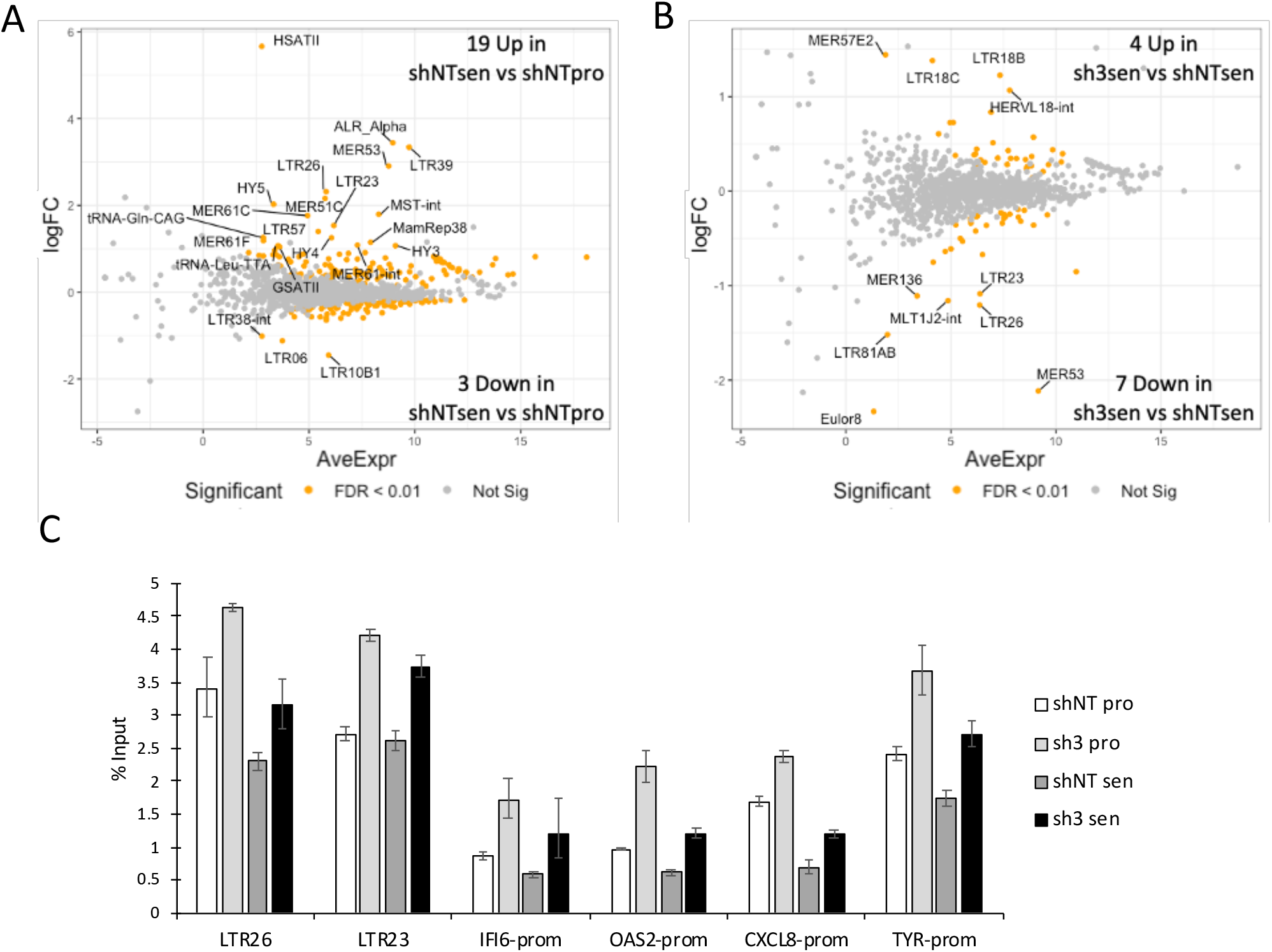
A subset of repeat RNAs are up-regulated in senescent versus proliferative sh-NT cells and down-regulated in senescent sh-H2AFJ versus senescent sh-NT cells. (**A**) Log2FC-Average Expression plots of repeat RNAs that are differentially expressed in senescent versus proliferative WI38hTERT/sh-NT cells. The majority of repeat RNAs are up-regulated in senescence. (**B**) LTR23, LTR26, and MER53 are up-regulated in senescent sh-NT cells and down-regulated in senescent sh-H2AFJ cells relative to sh-NT cells. (**C**) H1.2 ChIP qPCR showing that H1 occupancy is decreased at LTR23, LTR26, and IFI6, OAS2, CXCL8, and TYR promoters in senescent versus proliferative sh-NT cells. H1 occupancy is increased at all of these sites in sh-H2AFJ cells versus sh-NT cells in both proliferation and senescence.

### Senescent H2A.J-KO MEFs show defects in Interferon-Stimulared Gene expression

An H2A.J knock-out mouse was prepared as described in the Materials and Methods (Supplementary Figures 1A–1C) and used to prepare mouse embryonic fibroblasts (MEFs). We further tested a role for H2A.J in ISG expression by analyzing the transcriptome of WT and H2A.J MEFs induced into senescence by etoposide (Supplementary Figure 1D and 1E). The transcriptomes of senescent WT and H2A.J-KO showed strong separation from proliferating MEFs, and a weaker separation distinguished WT and H2A.J-KO MEFs (Supplementary Figure 4A). Strikingly, gene set enrichment analysis indicated highly significant defects in Interferon Response Gene Expression in the H2A.J-KO MEFs in senescence with significant down-regulation in senescent H2A.J-KO cells of a series of oligoadenylate synthase genes (Oas1g, Oas1a, Oasl1, Oas2, Oasl2) and several ISGs (Supplementary Figure 4C and 4D and Supplementary Table 7). Thus, H2A.J also contributes to ISG expression in the heterologous context of senescent MEFs.

### STAT1 activation is inhibited in senescent fibroblasts depleted for H2A.J

ISGs are transcriptionally activated by STAT and IRF transcription factors (43). STAT1, STAT2, IRF1, IRF2, and IRF7 genes were all transcriptionally up-regulated in senescent WI-38hTERT/sh-NT cells and down-regulated in senescent sh-H2AFJ cells, with STAT1 and IRF7 showing the greatest differential effect (Figure 6A). This deficit in STAT and IRF expression in the H2A.J-ko cells likely contributes to their lower level of ISG expression. Transcriptional activation of ISGs often involves phosphorylation and nuclear translocation of STAT transcription factors. Our transciptomic analysis indicated that STAT1 was the most highly transcribed STAT factor in WI-38 fibroblasts and its expression was activated 6-fold in senescence (Supplementary Table 6). Western blot and IF experiments showed that STAT1 levels, its phosphorylation, and its nuclear translocation were increased during etoposide-induced senescence, and all of these events were reduced in cells depleted for H2A.J (Figure 6B, 6C and 6D). Thus, H2A.J is required for maximal activation of the STAT1 pathway in senescence.

**Figure 6.**
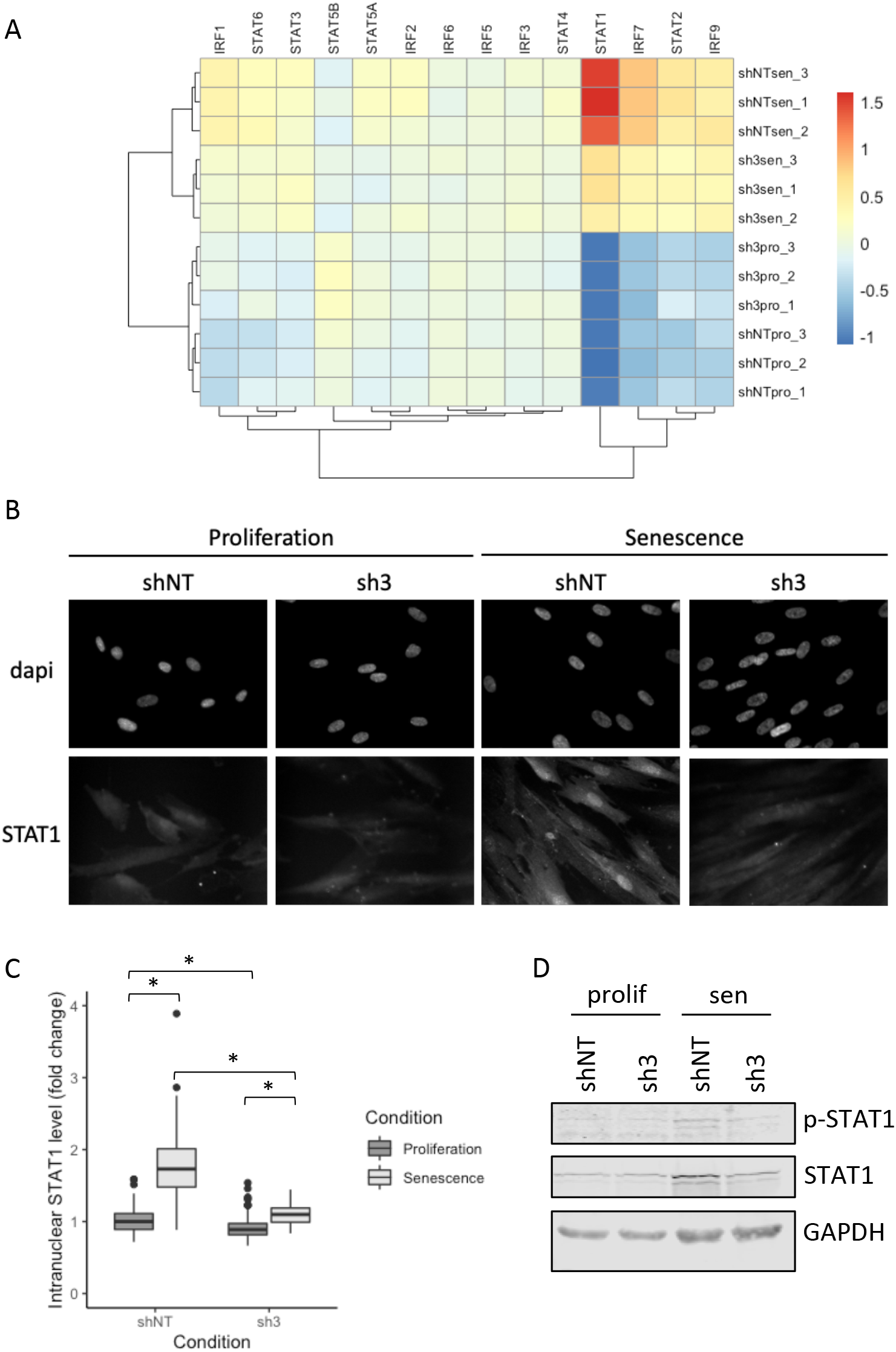
STAT1 activation in senescence is inhibited by H2A.J depletion. (**A**) Heat map of STAT and IRF RNA levels in sh-NT and sh3-H2AFJ cells in proliferation and senescence showing underexpression of several STAT and IRF genes in senescent sh3-H2AFJ cells relative to sh-NT cells. STAT1 is the most highly expressed STAT factor in senescent WI-38 fibroblasts. (**B**) IF images of STAT1 showing an increased intranuclear accumulation in senescent relative to proliferating sh-NT cells and a decreased signal in senescent sh-H2AFJ cells. (**C**) Quantification of the nuclear STAT1 signals for 3 independent IF experiments. Asterisk: p < 0.05 for a 2-sided t-test. Westen blot showing an increase of STAT1 and phospho-STAT1 in senescent relative to proliferating sh-NT cells and a decreased quantity in senescent sh3-H2AFJ cells.

### The Conserved H2A.J-Specific Val-11 and Ser-124 are implicated in its transcriptional activity

H2A.J differs from canonical H2A only by a valine at position 11 instead of alanine, and the 7 C-terminal amino acids containing a potential minimal phosphorylation site SQ for DNA-damage response kinases (Figure 7A). We previously showed that this C-terminal sequence is important for the transcriptional activity of H2A.J (11). To test the functional importance of these H2A.J-specific sequences, we mutated Val-11 to Ala as is found in all canonical H2A sequences, and we mutated Ser-123 to either Glu to mimic a phospho-serine residue or to Ala to prevent phosphorylation. We also substituted the C-terminus of H2A.J with the C-terminus of H2A. These mutants, WT-H2A.J and canonical H2A-type1 were ectopically expressed in proliferating fibroblasts, and their microarray transcriptomes were compared to that of proliferating and senescent fibroblasts without ectopic histone expression. Genome-wide transcriptome analysis indicated that senescent fibroblasts clustered distinctly from proliferating fibroblasts, and proliferating fibroblasts expressing the H2A.J-V11A and H2A.J-S123E mutants clustered distinctly from fibroblasts expressing the other H2A.J mutants, WT-H2A.J, and H2A (Figure 7B and Supplementary Table 8). Hallmark gene set enrichment analysis of the transcriptomes of fibroblasts expressing H2A.J-V11A or H2A.J-S123E versus control proliferating fibroblasts indicated that they showed the same highly significant enrichment for the Epithelial-Mesenchyme Transition, TNF-Alpha Signaling Via NF-kB, and Inflammatory Response gene sets (Figure 7C). Notable inflammatory genes including IL1A, IL1B, IL6, CXCL8, and CCL2 are contained in these gene sets and are often induced in senescence as part of the senescence-associated secretory phenotype (44). Heat maps showed that the H2A.J-V11A and H2A.J-S123E mutants were particularly apt at activating the expression of these inflammatory genes in proliferating fibroblasts (Figure 7D). RT-qPCR experiments confirmed these results for selected inflammatory genes (Supplementary Figure 5). Val11 thus appears to restrict the potential of H2A.J to promote inflammatory gene expression. The phospho-mimetic H2A.J-S123E mutant also hyperactivated inflammatory gene expression relative to WT-H2A.J or the non-phosphorylatable mutant H2A.J-S123A, suggesting that phosphorylation of S123 might be involved in its transcriptional activity.

**Figure 7.**
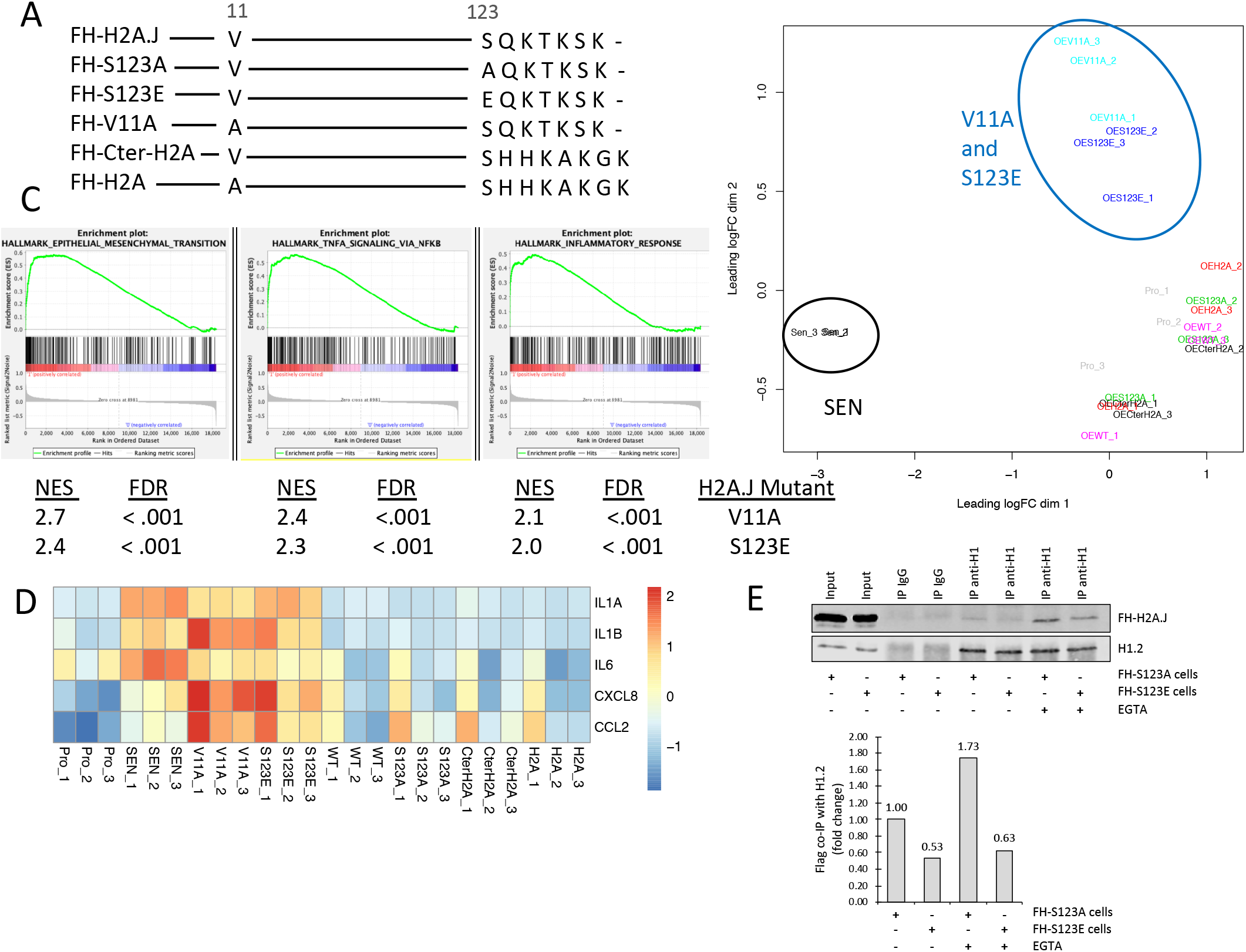
H2A.J-specific residues Val-11 and Ser-123 are functionally important. (**A**) Schematic showing H2A.J-specific residues, and mutants that were expressed in proliferating fibroblasts to examine their effect on gene expression. (**B**) MDS plot showing 3 distinct transcriptome clusters: senescent fibroblasts (SEN), H2A.J-V11A with H2A.J-S123E, and a final cluster of control proliferating fibroblasts and the same cells overexpressing the remaining mutants or canonical H2A. (**C**) GSEA showing highly significant enrichment of Hallmark Epithelial-Mesenchymal Transition, TNFA Signaling via NFkB, and Inflammatory Response gene sets for proliferating fibroblasts expressing H2A.J-V11A or H2A.J-S123E mutants. The False Discovery Rate (FDR) q-values and Normalized Enrichment Scores (NES) are shown below the enrichment plots. (**D**) Heat map of RNA levels for 6 inflammatory genes showing increased expression in proliferating fibroblasts expressing the H2A.J-V11A or H2A.J-S123E mutants. (**E**) The H2A.J-S123E mutant co-IPs less well with H1.2 compared to H2A.J-S123A.

H2A.J-S123 is expected to be in proximity to nucleosomal DNA (45, 46) and to the interaction site of H2A.J with H1 (8, 9), so phosphorylation of S123 might impact both of these interactions. We found that H2A.J-S123E co-immunoprecipitated less efficiently with H1.2 compared to H2A.J-S123A (Figure 7E), suggesting that phosphorylation of H2A.J-S123 would further weaken association of H2A.J with H1. We did not detect phosphorylation of H2A.J-S123 in senescent human fibroblasts by mass spectrometry. Since H2A.J is highly expressed in the liver and kidney of mice (11), we tested for phosphorylation of H2A.J 30 minutes after irradiating mice with 50 Gy of ionizing irradiation, and we observed S123 phosphorylation at an estimated level of about 1% of all H2A.J molecules (Supplementary Figure 6). In contrast, in the same sample, more than 50% of all H2A.X molecules were phosphorylated at its C-terminal SQE motif. Thus, in response to ionizing irradiation, H2A.X is phosphorylated much more efficiently than H2A.J. A more sensitive detection method is necessary to determine whether H2A.J is phosphorylated at low levels in senescent fibroblasts. Overall, our mutational analysis indicates that the conserved Val11 and Ser-123 are functionally important for H2A.J transcriptional activity.

## DISCUSSION

Our previous work indicated that accumulation of H2A.J promoted the expression of inflammatory genes in senescent WI-38 fibroblasts with persistent DNA damage (11). Furthermore, over-expression of H2A.J in proliferating fibroblasts increased inflammatory gene expression, and in particular, Interferon-Stimulated Genes (ISGs). Derepression of repeated DNA sequences has been reported in some senescent cells (42, 47), and we confirmed this finding for our WI-38 fibroblasts induced into senescence by treatment with the DNA-damaging agent etoposide. Repeat RNAs can induce the expression of ISGs (7, 42). In this work, we show that some repeat RNAs are expressed at lower levels in senescent WI-38 cells depleted for H2A.J, which may contribute to the decreased ISG expression observed in these cells relative to sh-NT control cells. We suggest that derepression of repeated DNA transcription in senescence is explained at least in part by the reported decrease in histone H1 levels in senescent cells (38). We show here that that accumulation of H2A.J in senescent cells contributes to decreasing the half-life of histone H1 by a weakened interaction of H2A.J with H1 compared to H2A with H1. We suggest that the stability of H1 is controlled by its association with chromatin and with histone chaperone complexes in the nucleosoluble compartment, and these associations are weakened when H2A.J levels increase in senescence leading to increased turnover of H1. The C-terminus of H2A was previously shown to interact with H1 (8, 9), so it is plausible that the divergent C-terminus of H2A.J relative to H2A could weaken its interaction with H1. H2A forms a heterodimeric complex with H2B rapidly after its synthesis (36, 37). In the nucleosoluble compartment of cells, we find H1 in gel filtration fractions containing H2A, H2B, and histone chaperones including NASP, NAP1L, and FACT. We thus suggest that nucleosoluble H1 interacts with H2A and H2B in complex with histone chaperones, but further work is necessary to precisely characterize these complexes. Interestingly, the association of H2A with H1 is stabilized in the presence of the calcium chelator EGTA. Calcium was previously shown to influence the secondary structure of H1 (35), and calcium levels are increased in senescent cells (48). We thus suggest that both increased H2A.J expression and increased calcium levels in senescent cells participate in destabilizing the interaction of H1 with chromatin thereby increasing the turnover of H1 in senescence. Further factors leading to H1 decreases in senescence include transcriptional repression of H1 genes, and the yet uncharacterized pathway responsible for H1 turnover (38). Decreased H1 levels in senescence would then contribute to the derepression of some repeated DNA sequences and consequent activation of ISG expression (Figure 8). This model is consistent with previous work indicating that H1 levels decrease in senescent WI-38 fibroblasts (38), and loss of H1 leads to transcriptional derepression of repeated DNA sequences and ISG expression (7). This model requires further testing. Unfortunately, previous work indicated that over-expressing H1 in senescent cells did not allow significant restoration of H1 levels (38). Thus, we need to better characterize the pathway leading to H1 proteolysis in senescence so that we can restore H1 levels and test its effect on ISG expression. The innate immunity proteins that recognize repeat RNAs in senescence must also be identified (49).

**Figure 8.**
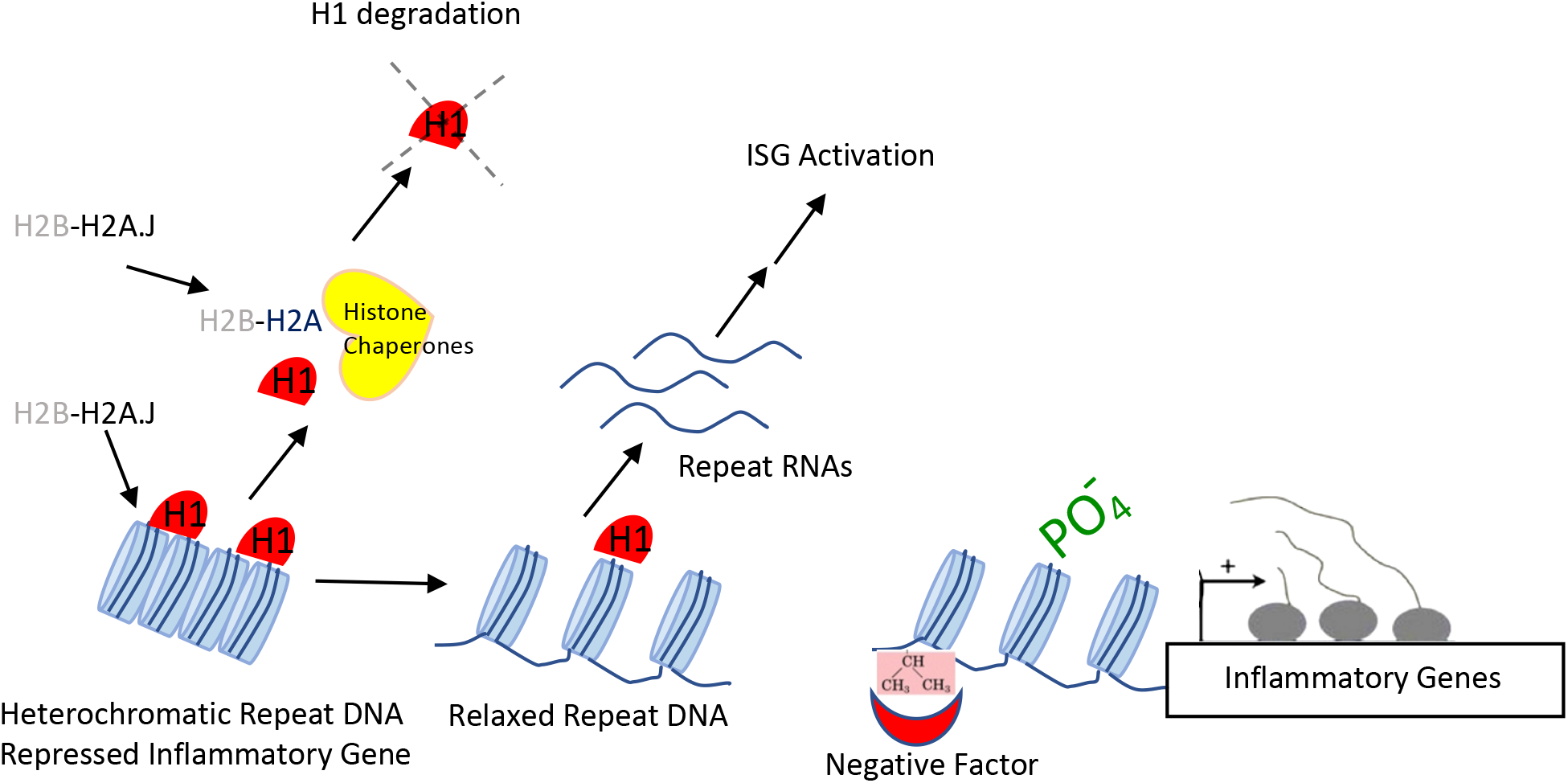
Speculative model summarizing H2A.J Function in Promoting Inflammatory Gene Expression. Accumulation of H2A.J in senescence competes with H2A for deposition in chromatin and association with H1 in nucleosoluble complexes. The weaker association of H2A.J for H1, and increased calcium in senescence, contributes to weaker association of H1 with chromatin and increased H1 turnover leading to increased transcription of some repeated DNA elements. Repeat RNAs would trigger innate immune pathways leading to STAT/IRF activation and ISG transcription. Phosphorylation of H2A.J-S123 in response to DNA damage may contribute to transcriptional activation of inflammatory genes by further weakening association of H2A.J with H1 and by decreasing the affinity of DNA with nucleosomes in the vicinity of inflammatory genes. H2A.J-Val11 moderates H2A.J transcriptional activity, so it may participate in recruiting a factor that limits inflammatory gene transcription.

Transcriptional activation of ISGs is mediated by STAT and IRF transcription factors (43). We found that STAT1, STAT2, IRF1, IRF2, and IRF7 mRNA levels increased in senescent WI-38 control cells, and this induction was inhibited in cells knocked-down for H2A.J. Consistently, nuclear STAT1 protein and phospho-STAT1 were reduced in senescent cells depleted for H2A.J. The reduced expression of STAT/IRF factors could explain the reduced expression of ISGs in H2A.J-depleted senescent cells.

Transcriptional activation of interferon (IFN) genes by STAT/IRF factors serves in an auto-activation loop to boost expression of ISGs (43). The IFNB1 gene was induced in senescent WI-38 cells, but its expression level was low (Supplementary Table 6). De Cecco et al. showed that IFN gene activation is a late event in the DNA-damage induced senescence of fibroblasts (42). We analyzed cells at 7 days after inducing senescence with etoposide, so we suggest that the induction of IFN gene expression was only just beginning at this early time point. These authors also showed that the RB1 transcriptional repressor contributes to repression of L1 repeated DNAs in senescence. It remains to be seen whether H1 and Rb contribute independently to repression of repeated DNA or whether they are part of the same pathway.

H2A.J differs from canonical H2A only by a substitution of valine for alanine at position 11, and in the last 7 amino acids containing an SQ motif in H2A.J that could potentially be phosphorylated on Ser-123. We tested the effect on inflammatory gene expression of ectopically expressing mutants of these conserved residues of H2A.J in proliferating fibroblasts relative to wild-type H2A.J and canonical H2A-type1. The H2A.J-V11A mutant increased expression of inflammatory genes to a greater extent than wild-type H2A.J, indicating that Val11 normally moderates the potential of H2A.J to promote inflammatory gene expression. Valine differs from alanine by the addition of 2 methyl groups on its side gene. The N-terminal tails of core histones protrude from the nucleosome surface such that they can interact with chromatin regulators. Methylation of histone N-terminal lysine and arginine residues can contribute to both activation or repression of transcription by forming binding surfaces for the recruitment of transcriptional regulators (50). We speculate that the dimethyl groups of H2A.J-Val11 form part of a binding site for a regulatory factor that limits its potential to promote inflammatory gene expression. We also mutated Ser-123 to Ala to block phosphorylation and to Glu to mimic phosphorylation. The H2A.J-S123E mutant hyper-activated inflammatory gene expression whereas the H2A.J-S123A mutant acted similarly to WT-H2A.J. Ser-123 is close to nucleosomal DNA (45, 46) and its phosphorylation could weaken the interaction of DNA with the nucleosome to facilitate transcription. Ser-123 is also close to the site of interaction of H2A with H1 (8, 9) and we found that the H2A.J-S123E mutant interacted less well with H1 compared to H2A.J which in turn interacts less well than H2A. Thus, H2A.J-S123 phosphorylation could potentially promote inflammatory gene expression by both loosening the association of DNA with the nucleosome and by weakening the interaction of H1 with chromatin. Mass spectrometry experiments revealed that H2A.J was phosphorylated at Ser-123 after irradiation, but at levels that were much lower than for H2A.X. Two factors may contribute to the lower phosphorylation of H2A.J by DNA damage response (DDR) kinases. First, the H2A.J-SQK sequence may be a less optimal substrate than the H2A.X-SQE motif for DDR kinases, and second, the longer C-terminal tail of H2A.X would lead to its greater extrusion from the nucleosome that would facilitate access of the DDR kinases to the SQ motif. We were unable to detect phosphorylation of H2A.J-Ser123 in senescent human fibroblasts. If H2A.J was phosphorylated at only a small number of sites near genes that it regulates, then we may have been unable to detect it with the reagents and methods currently at our disposal. High affinity anti-phospho-S123-H2A.J antibodies will be necessary to test for low-level phosphorylation of H2A.J at specific chromatin sites. Our current work has demonstrated the functional importance of conserved H2A.J N-terminal and C-terminal residues and it has suggested potential mechanisms for their action in terms of interactions with H1 and other negative transcriptional regulators, and the wrapping of DNA around nucleosomes (Figure 8).

## SUPPLEMENTARY DATA

**Supplementary Table 1**. DNA sequences of synthetic DNA fragments encoding Flag-HA (FH) tagged H2A.J, the indicated H2A.J mutants, and H2A. These fragments were cloned into the Age1/MluI sites of lentiviral pTRIPz to place the coding sequences under the control of the tet-ON promoter.

**Supplementary Table 2**. Sequence of primers used for qPCR

**Supplementary Table 3**. Tandem mass spectrometry analysis of proteins co-purifying with Flag-HA-H2A.J from the nucleosoluble fraction of HeLa S3 cells.

**Supplementary Table 4**. Tandem mass spectrometry analysis of proteins co-purifying with H1 and H2A after Superose 6 gel filtration of T47D nucleosoluble extracts.

**Supplementary Table 5**. Levels of small RNAs (TPM) and differentially-expressed genes for WI-38hTERT sh-NT and sh3-H2AFJ cells in proliferation and etoposide-induced senescence.

**Supplementary Table 6**. Levels of total RNAs (TPM) after rRNA depletion, and differentially-expressed genes, for WI-38hTERT sh-NT and sh3-H2AFJ cells in etoposide-induced senescence and for sh-NT cells in senescence versus proliferation.

**Supplementary Table 7**. RNA levels (TPM) and differentially-expressed genes for WT and H2A.J-KO MEFs in proliferation and etoposide-induced senescence.

**Supplementary Table 8**. Gene expression levels and differentially expressed genes from Illumina Bead Chip microarrays of WI-38hTERT cells in proliferation and etoposide-induced senescence, and of proliferating WI-38hTERT cells over-expressing WT-H2A.J, H2A.J mutants, and canonical H2A-type1.

Supplementary Data are available at NAR online.

## ACKNOWLEDGEMENT

This work has benefited from the facilities and the expertise of the High-throughput Sequencing Platform and the SICaPS Proteomic platform of the I2BC (Institute for Integrative Biology of the Cell), CEA, CNRS, Univ. Paris-Sud, Université Paris-Saclay, 91198, Gif-sur-Yvette cedex, France.

## TABLE AND FIGURES LEGENDS

**Supplementary Table 2**. Sequence of primers used for qPCR

**Supplementary Figure 1.**
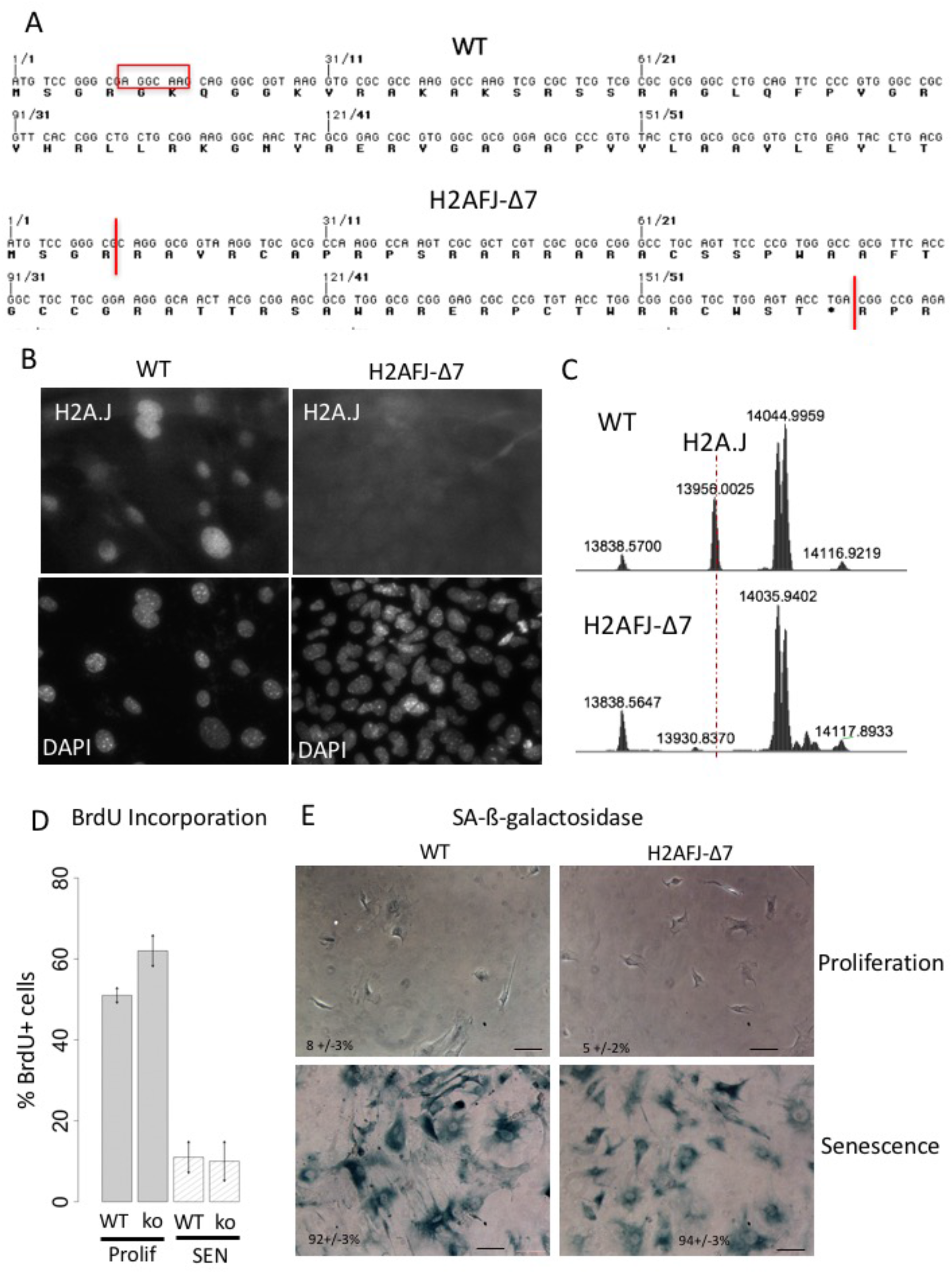
H2A.J-KO mouse and MEFs. (**A**) Nucleotide and encoded amino acid sequences of the beginning of the murine H2AFJ-WT and H2AFJΔ7 TALEN-mediated 7 bp deletion mutant showing the frameshift and premature stop codon introduced by the mutation. (**B**) Immunofluorescence assay with anti-H2A.J antibodies showing the absence of H2A.J expression in H2AFJΔ7-KO MEFs relative to WT MEFs. (**C**) Top-down mass spectrometry analysis of H2A type1 histones acid-extracted from the kidneys of 3 month old WT and H2AFJΔ7-KO mice showing the absence of H2A.J in the H2AFJΔ7-KO kidneys. Mass spectrometry was performed as previously described (Contrepois et al., 2017). (**D, E**) Induction of senescence after treating WT and H2A.J-ko MEFs with 2.5 μM etoposide. (**D**) BrdU incorporation assay showing the proliferative arrest induced by etoposide treatment (SEN). (**E**) Phase contrast images of WT and H2A.J-KO MEFs stained for SA-ß-galactosidase activity and percent positive cells.

**Supplementary Figure 2:**
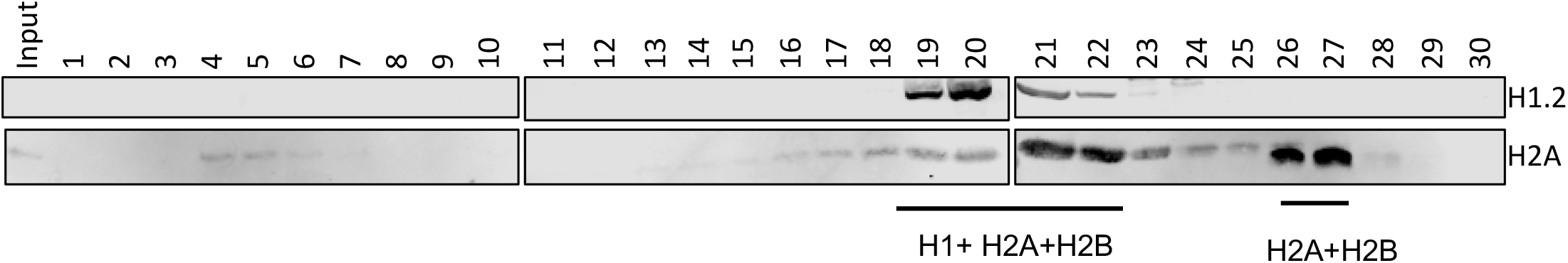
Superose 6 gel filtration of nucleoplasmic extracts of T47D cells. Shown are Western blots to identify the elution fractions containing histones H1.2 and H2A.

**Supplementary Figure 3.**
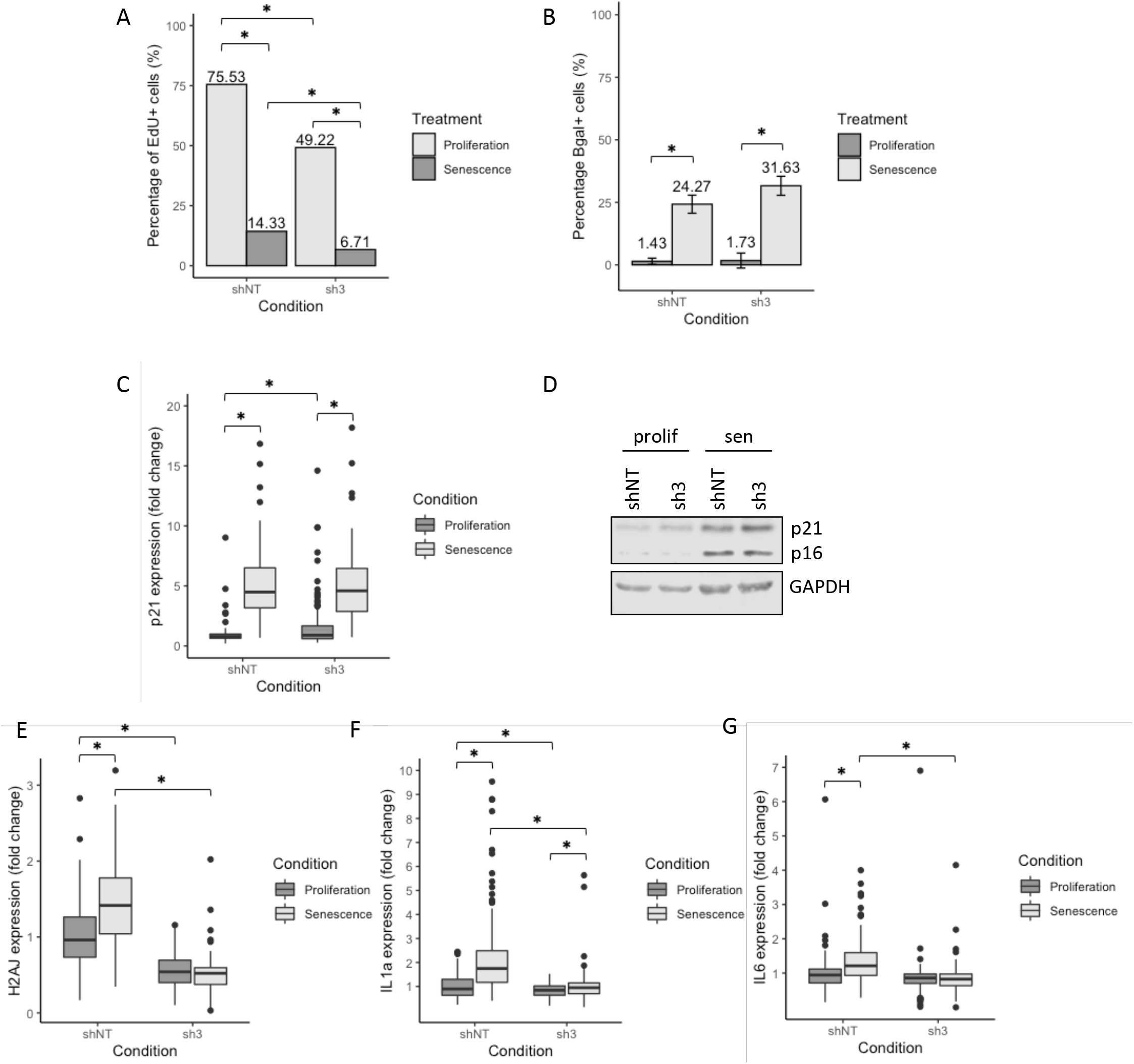
Assays demonstrating induction of senescence with activation of inflammatory gene expression for sh-NT and sh3-H2AFJ WI-38hTERT fibroblasts in proliferation and after treatment with 100 μM etoposide for 2 days followed by 7 days in medium without etoposide. (**A**) Percentage of EdU-positive cells to assay proliferation. (**B**) Percentage of SA-ß-galactosidase positive cells. (**C**) Quantification of p21 levels by an immunofluorescence assay. (**D**) p16 and p21 Western blots. One representative experiment of 3. Quantification of H2A.J (**E**), IF1A (**F**), and IL6 (**G**) levels by an immunofluorescence assay. Quantifications were done on 3 independent experiments. Asterisks: p < 0.05 by 2-sided t-tests.

**Supplementary Figure 4.**
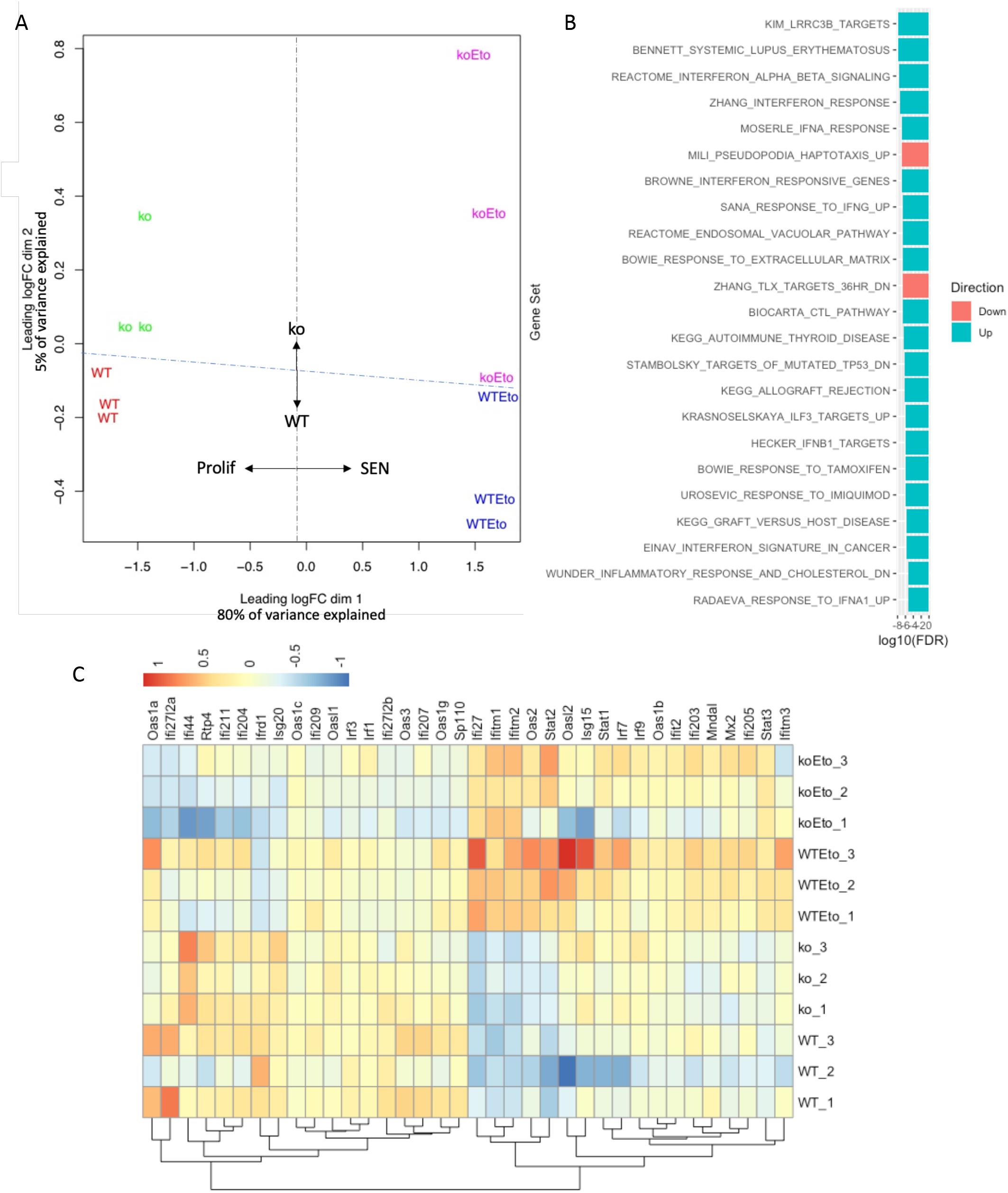
Senescent H2A.J-KO MEFs show defects in ISG expression. (**A**) Limma (Ritchie et al, 2015) multidimensional scaling plot (MDS) of the transciptomes of WT and H2AFJ-KO MEFs in proliferation and senescence showing strong variation in the transcriptomes of cells in senescence versus proliferation, and a weaker variation distinguishing WT and H2A.J-KO MEFs. (**B**) GSEA using the Broad Institute c2 curated gene sets on the transcriptomes of WT and H2AFJ-KO MEFs in senescence versus proliferation showing enrichment for up-regulation of many Interferon and Immune Response gene sets. (**C**) Heat map of genes within the Interferon Response gene sets.

**Supplementary Figure 5.**
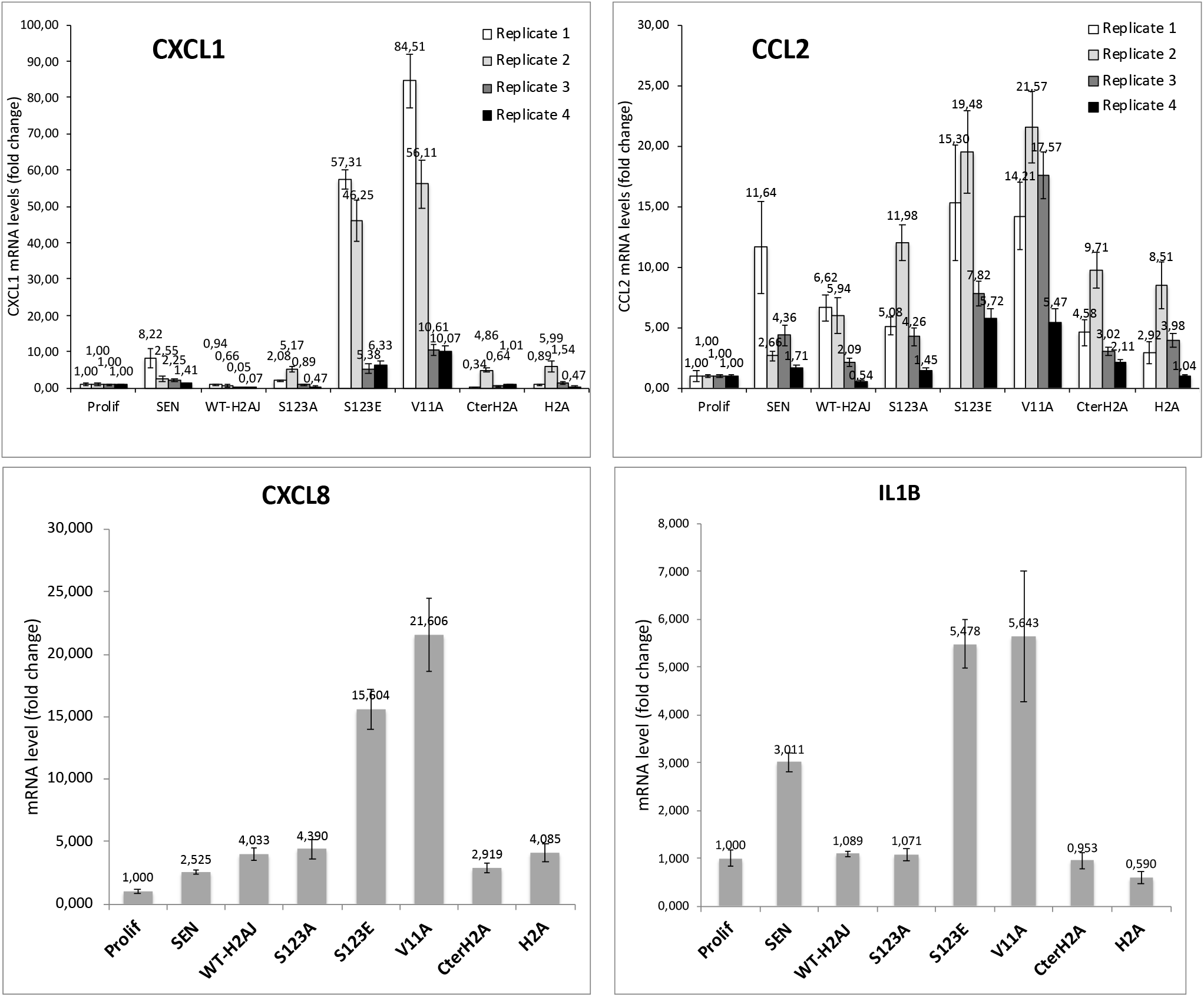
RT-qPCR of CXCL1, CCL2, IL1B and CXCL8 mRNA levels in WI-38hTERT cells in proliferation and etoposide-induced senescence, and in proliferating cells overexpressing WT-H2A.J, the indicated H2A.J mutants, and canonical H2A. 4 biological replicates are shown for CXCL1 and CCL2. Although the magnitude of gene induction varied in the different replicates, expression of H2A.J-S123E and H2A.J-V11A always led to the highest levels of CXCL1 and CCL2 induction. Only one replicate was done for CXCL8 and IL1B.

**Supplementary Figure 6.**
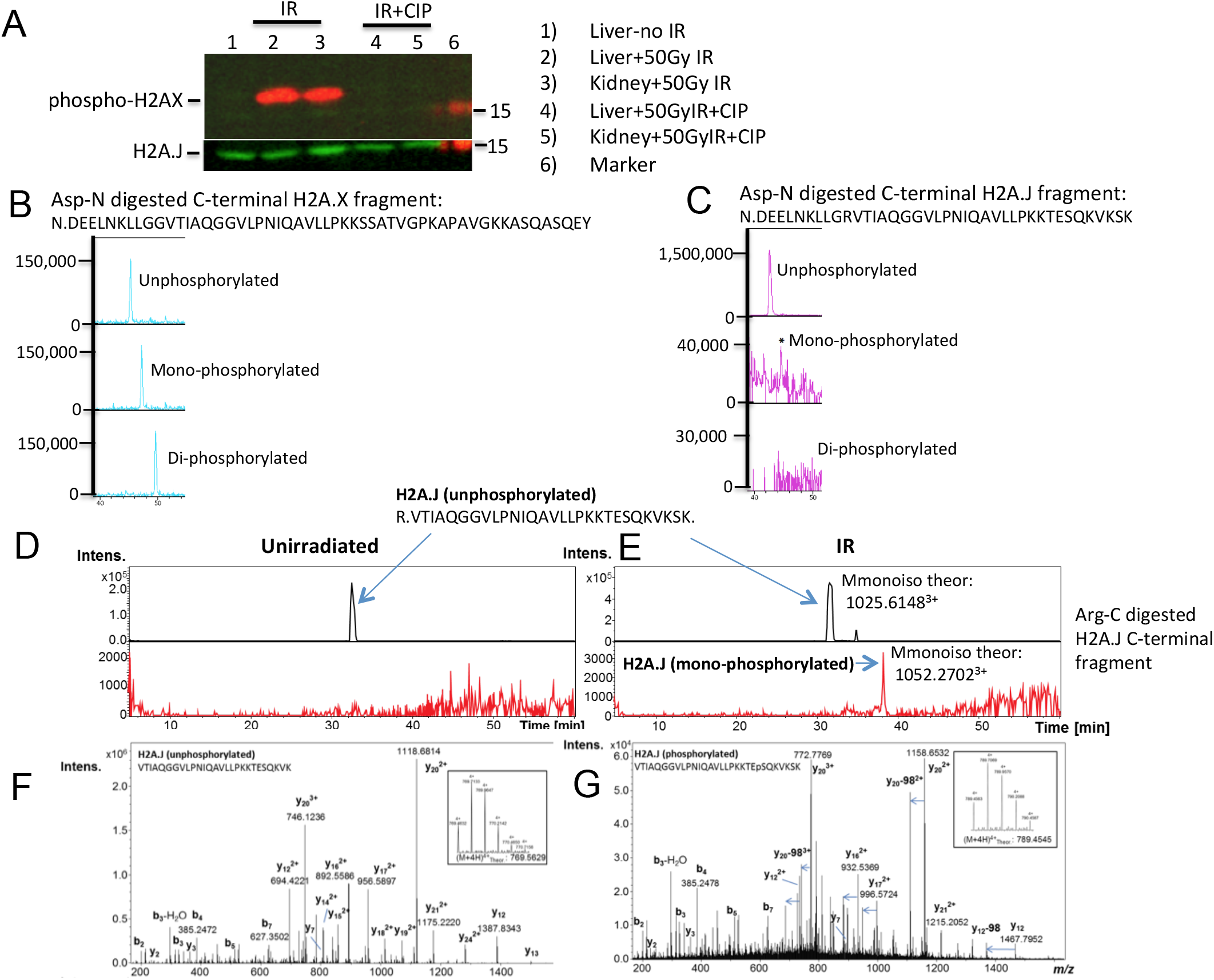
H2A.J is phosphorylated at S123 in mouse kidneys 30 minutes after 50 Gy irradiation. (**A**) Western blot of H2A.J and phospho-H2A.X in acid-extracted histones prepared from the liver and kidney of a mouse 30 minutes post-irradiation with 50 Gy of ionizing irradiation (IR) or a control non-irradiated mouse. An aliquot of the histones was also treated with calf intestinal phosphatase (CIP). The Western blot shows induced phosphorylation of H2A.X after IR and constant levels of H2A.J. (**B,C**) Tandem LC-MS/MS of Asp-N digested histones showing spectral counts for the C-terminal unphosphorylated and phosphorylated C-terminal H2A.X and H2A.J peptides. (**D,E**) Tandem LC-MS/MS of Arg-C digested histones showing spectral counts for the C-terminal unphosphorylated and phosphorylated C-terminal H2A.J peptides. (**F,G**) Fragmentation patterns of the Arg-C unphosphorylated and phosphorylated C-terminal H2A.J peptides allowing manual identification of these peptides and phosphorylation of H2A.J on Ser-123. Of note, MS/MS spectra of phospho-peptides typically contain an intense neutral loss peak that resides at 98 Da lower than some ions, representing the loss of H_3_PO_4_ (indicated in the figure as “-98” and with a blue arrow).

**Supplementary Table 1.**
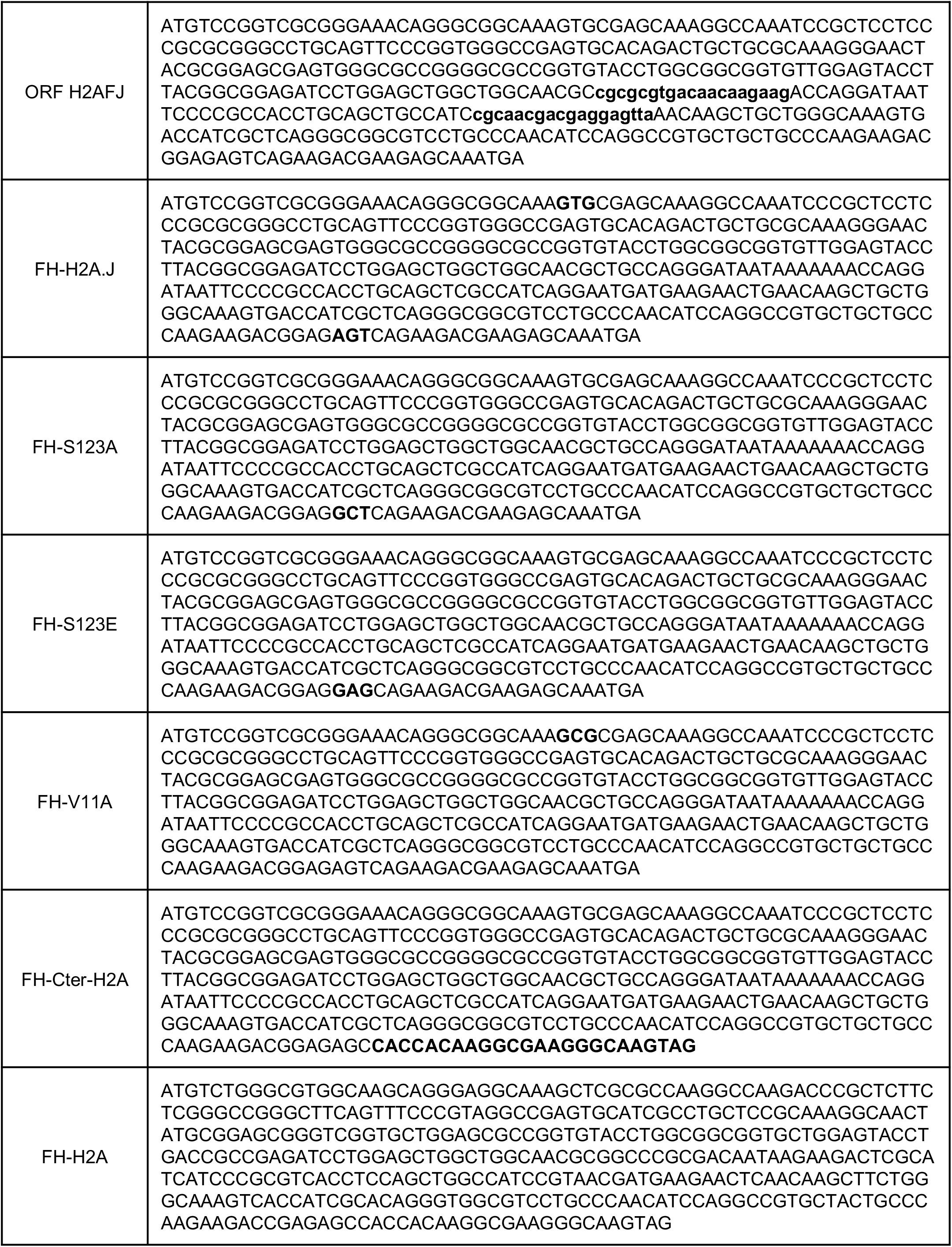
DNA sequences of synthetic DNA fragments encoding Flag-HA (FH) tagged H2A.J, the indicated H2A.J mutants, and H2A. These fragments were cloned into the Age1/MluI sites of lentiviral pTRIPz to place the coding sequences under the control of the tet-ON promoter.

**Supplementary Table 2.**
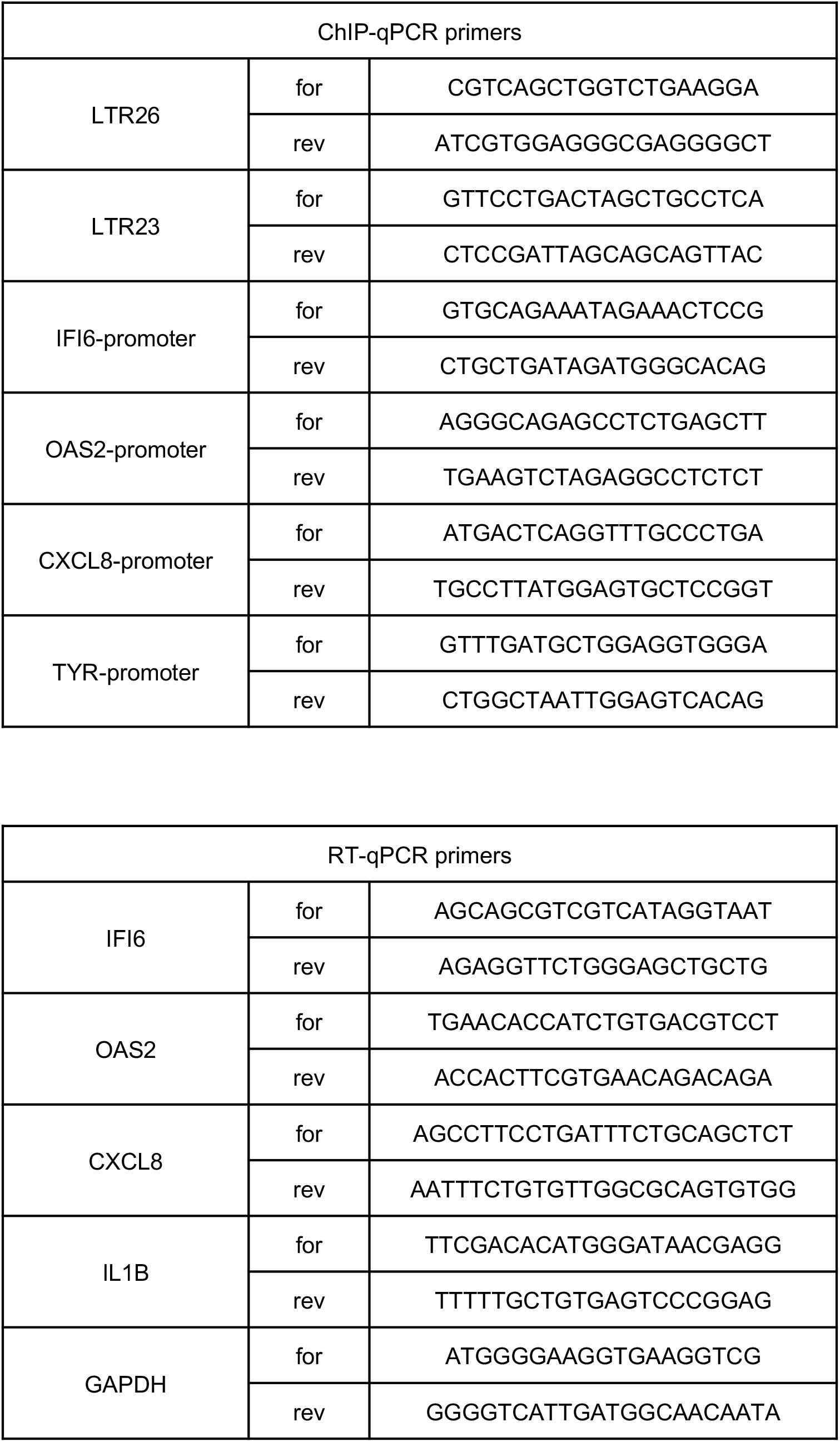
Sequence of primers used for qPCR.

**Supplementary Table 3.** Tandem mass spectrometry analysis of proteins co-purifying with Flag-HAH2A. J from the nucleosoluble fraction of HeLa S3 cells.

**Supplementary Table 4.** Tandem mass spectrometry analysis of proteins co-purifying with H1 and H2A after Superose 6 gel filtration of T47D nucleosoluble extracts.

## REFERENCES

1. Cutter,A.R.R. and Hayes,J.J.J. (2015) A brief review of nucleosome structure. FEBS Lett., 10.1016/j.febslet.2015.05.016.

2. Marzluff,W.F.F. and Koreski,K.P.P. (2017) Birth and Death of Histone mRNAs. Trends Genet., 10.1016/j.tig.2017.07.014.

3. Skene,P.J. and Henikoff,S. (2013) Histone variants in pluripotency and disease. Development, 140, 2513–24.

4. Shi,L., Wen,H. and Shi,X. (2017) The Histone Variant H3.3 in Transcriptional Regulation and Human Disease. J. Mol. Biol., 10.1016/j.jmb.2016.11.019.

5. Bonner,W.M., Redon,C.E., Dickey,J.S., Nakamura,A.J., Sedelnikova,O.A., Solier,S. and Pommier,Y. (2008) γH2AX and cancer. Nat. Rev. Cancer, 8, 957–967.

6. Hergeth,S.P.P. and Schneider,R. (2015) The H1 linker histones: multifunctional proteins beyond the nucleosomal core particle. EMBO Rep., 10.15252/embr.201540749.

7. Izquierdo-Bouldstridge,A., Bustillos,A., Bonet-Costa,C., Aribau-Miralbés,P., García-Gomis,D., Dabad,M., Esteve-Codina,A., Pascual-Reguant,L., Peiró,S., Esteller,M., et al. (2017) Histone H1 depletion triggers an interferon response in cancer cells via activation of heterochromatic repeats. Nucleic Acids Res., 10.1093/nar/gkx746.

8. Shukla,M.S., Syed,S.H., Goutte-Gattat,D., Richard,J.L.C., Montel,F., Hamiche,A., Travers,A., Faivre-Moskalenko,C., Bednar,J., Hayes,J.J., et al. (2011) The docking domain of histone H2A is required for H1 binding and RSC-mediated nucleosome remodeling. Nucleic Acids Res., 39, 2559–2570.

9. Vogler,C., Huber,C., Waldmann,T., Ettig,R., Braun,L., Izzo,A., Daujat,S., Chassignet,I., Lopez-Contreras,A.J.J., Fernandez-Capetillo,O., et al. (2010) Histone H2A C-terminus regulates chromatin dynamics, remodeling, and histone H1 binding. PLoS Genet., 10.1371/journal.pgen.1001234.

10. Nishida,H., Suzuki,T., Tomaru,Y. and Hayashizaki,Y. (2005) A novel replication-independent histone H2a gene in mouse. BMC Genet., 6, 10.

11. Contrepois,K., Coudereau,C., Benayoun,B., Schuler,N., Courbeyrette,R., Thuret,J.-Y., Ma,Z., Derbois,C., Nevers,M., Snyder,M., et al. (2017) A novel histone variant H2A. J accumulates in senescent human cells with persistent DNA damage and promotes inflammatory gene expression. Nat. Commun., 8, 1–15.

12. Capell,B.C., Drake,A.M., Zhu,J., Shah,P.P., Dou,Z., Dorsey,J., Simola,D.F., Donahue,G., Sammons,M., Rai,T.S., et al. (2016) Mll1 is essential for the senescenceassociated secretory phenotype. Genes Dev., 30, 321–336.

13. Barde,I., Salmon,P. and Trono,D. (2010) Production and titration of lentiviral vectors. Curr. Protoc. Neurosci., 10.1002/0471142301.ns0100s37.

14. Durkin,M., Qian,X., Popescu,N. and Lowy,D. (2013) Isolation of Mouse Embryo Fibroblasts. BIO-PROTOCOL, 10.21769/bioprotoc.908.

15. Kunieda,T., Xian,M., Kobayashi,E., Imamichi,T., Moriwaki,K. and Toyoda,Y. (1992) Sexing of mouse preimplantation embryos by detection of Y chromosome-specific sequences using polymerase chain reaction. Biol. Reprod., 10.1095/biolreprod46.4.692.

16. Goyallon,A., Cholet,S., Chapelle,M., Junot,C. and Fenaille,F. (2015) Evaluation of a combined glycomics and glycoproteomics approach for studying the major glycoproteins present in biofluids: Application to cerebrospinal fluid. Rapid Commun. Mass Spectrom., 10.1002/rcm.7125.

17. Obri,A., Ouararhni,K., Papin,C., Diebold,M.L.L., Padmanabhan,K., Marek,M., Stoll,I., Roy,L., Reilly,P.T.T., Mak,T.W.W., et al. (2014) ANP32E is a histone chaperone that removes H2A.Z from chromatin. Nature, 10.1038/nature12922.

18. Henikoff,S., Henikoff,J.G., Sakai,A., Loeb,G.B. and Ahmad,K. (2009) Genome-wide profiling of salt fractions maps physical properties of chromatin. Genome Res., 19, 460–469.

19. Dimri,G.P., Lee,X., Basile,G., Acosta,M., Scott,G., Roskelley,C., Medrano,E.E., Linskens,M., Rubelj,I., Pereira-Smith,O., et al. (1995) A biomarker that identifies senescent human cells in culture and in aging skin in vivo. Proc Natl Acad Sci U S A, 92, 9363–9367.

20. Martin,M. (2011) Cutadapt removes adapter sequences from high-throughput sequencing reads. EMBnet.journal, 17, 10.

21. Andrews,S. (2010) FastQC: A quality control tool for high throughput sequence data. http://www.bioinformatics.babraham.ac.uk/projects/fastqc/, citeulike-article-id:11583827.

22. Li,H. (2013) Aligning sequence reads, clone sequences and assembly contigs with BWA-MEM.

23. Langmead,B. and Salzberg,S.L. (2012) Fast gapped-read alignment with Bowtie 2. Nat. Methods, 10.1038/nmeth.1923.

24. Feng,J., Liu,T., Qin,B., Zhang,Y. and Liu,X.S. (2012) Identifying ChIP-seq enrichment using MACS. Nat. Protoc., 10.1038/nprot.2012.101.

25. Yu,G., Wang,L.G. and He,Q.Y. (2015) ChIP seeker: An R/Bioconductor package for ChIP peak annotation, comparison and visualization. Bioinformatics, 10.1093/bioinformatics/btv145.

26. Yu,G. and He,Q.Y. (2016) ReactomePA: An R/Bioconductor package for reactome pathway analysis and visualization. Mol. Biosyst., 10.1039/c5mb00663e.

27. Patro,R., Duggal,G., Love,M.I., Irizarry,R.A. and Kingsford,C. (2017) Salmon provides fast and bias-aware quantification of transcript expression. Nat. Methods, 10.1038/nmeth.4197.

28. Love,M.I., Soneson,C., Hickey,P.F., Johnson,L.K., Tessa Pierce,N., Shepherd,L., Morgan,M. and Patro,R. (2020) Tximeta: Reference sequence checksums for provenance identification in RNA-seq. PLoS Comput. Biol., 10.1371/journal.pcbi.1007664.

29. Love,M.I., Anders,S. and Huber,W. (2014) Differential analysis of count data-the DESeq2 package.

30. Anders,S., McCarthy,D.J., Chen,Y., Okoniewski,M., Smyth,G.K., Huber,W. and Robinson,M.D. (2013) Count-based differential expression analysis of RNA sequencing data using R and Bioconductor. Nat. Protoc., 10.1038/nprot.2013.099.

31. Ritchie,M.E., Phipson,B., Wu,D., Hu,Y., Law,C.W., Shi,W. and Smyth,G.K. (2015) Limma powers differential expression analyses for RNA-sequencing and microarray studies. Nucleic Acids Res., 43, e47.

32. Law,C.W., Chen,Y., Shi,W. and Smyth,G.K. (2014) Voom: Precision weights unlock linear model analysis tools for RNA-seq read counts. Genome Biol., 10.1186/gb-2014-15-2-r29.

33. Wu,D. and Smyth,G.K. (2012) Camera: A competitive gene set test accounting for inter-gene correlation. Nucleic Acids Res., 10.1093/nar/gks461.

34. Subramanian,A., Tamayo,P., Mootha,V.K., Mukherjee,S., Ebert,B.L., Gillette,M.A., Paulovich,A., Pomeroy,S.L., Golub,T.R., Lander,E.S., et al. (2005) Gene set enrichment analysis: a knowledge-based approach for interpreting genome-wide expression profiles. Proc Natl Acad Sci U S A, 102, 15545–15550.

35. Tarkka,T., Oikarinen,J. and Grundström,T. (1997) Nucleotide and calcium-induced conformational changes in histone H1. FEBS Lett., 406, 56–60.

36. Mosammaparast,N., Ewart,C.S. and Pemberton,L.F. (2002) A role for nucleosome assembly protein 1 in the nuclear transport of histones H2A and H2B. EMBO J., 10.1093/emboj/cdf647.

37. Chang,L., Loranger,S.S., Mizzen,C., Ernst,S.G., Allis,C.D. and Annunziato,A.T. (1997) Histones in transit: Cytosolic histone complexes and diacetylation of H4 during nucleosome assembly in human cells. Biochemistry, 10.1021/bi962069i.

38. Funayama,R., Saito,M., Tanobe,H. and Ishikawa,F. (2006) Loss of linker histone H1 in cellular senescence. J Cell Biol, 175, 869–880.

39. Bhaumik,D., Scott,G.K.K., Schokrpur,S., Patil,C.K.K., Orjalo,A.V. V., Rodier,F., Lithgow,G.J.J. and Campisi,J. (2009) MicroRNAs miR-146a/b negatively modulate the senescence-associated inflammatory mediators IL-6 and IL-8. Aging (Albany. NY)., 10.18632/aging.100042.

40. Taganov,K.D.D., Boldin,M.P.P., Chang,K.J.J. and Baltimore,D. (2006) NF-κB-dependent induction of microRNA miR-146, an inhibitor targeted to signaling proteins of innate immune responses. Proc. Natl. Acad. Sci. U. S. A., 10.1073/pnas.0605298103.

41. De Cecco,M., Criscione,S.W.W., Peckham,E.J.J., Hillenmeyer,S., Hamm,E.A.A., Manivannan,J., Peterson,A.L.L., Kreiling,J.A.A., Neretti,N. and Sedivy,J.M.M. (2013) Genomes of replicatively senescent cells undergo global epigenetic changes leading to gene silencing and activation of transposable elements. Aging Cell, 10.1111/acel.12047.

42. De Cecco,M., Ito,T., Petrashen,A.P.P., Elias,A.E.E., Skvir,N.J.J., Criscione,S.W.W., Caligiana,A., Brocculi,G., Adney,E.M.M., Boeke,J.D.D., et al. (2019) L1 drives IFN in senescent cells and promotes age-associated inflammation. Nature, 10.1038/s41586-018-0784-9.

43. Michalska,A., Blaszczyk,K., Wesoly,J. and Bluyssen,H.A.R.A.R. (2018) A positive feedback amplifier circuit that regulates interferon (IFN)-stimulated gene expression and controls type I and type II IFN responses. Front. Immunol., 10.3389/fimmu.2018.01135.

44. Coppé,J.P., Patil,C.K., Rodier,F., Sun,Y., Muñoz,D.P., Goldstein,J., Nelson,P.S., Desprez,P.Y. and Campisi,J. (2008) Senescence-associated secretory phenotypes reveal cell-nonautonomous functions of oncogenic RAS and the p53 tumor suppressor. PLoS Biol., 10.1371/journal.pbio.0060301.

45. Li,Z. and Kono,H. (2016) Distinct Roles of Histone H3 and H2A Tails in Nucleosome Stability. Sci. Rep., 10.1038/srep31437.

46. Tanaka,H., Sato,S., Koyama,M., Kujirai,T. and Kurumizaka,H. (2020) Biochemical and structural analyses of the nucleosome containing human histone H2A.J. J. Biochem., 10.1093/jb/mvz109.

47. Sidler,C., Woycicki,R., Li,D., Wang,B., Kovalchuk,I. and Kovalchuk,O. (2014) A role for SUV39H1-mediated H3K9 trimethylation in the control of genome stability and senescence in WI38 human diploid lung fibroblasts. Aging (Albany. NY)., 10.18632/aging.100678.

48. Martin,N. and Bernard,D. (2018) Calcium signaling and cellular senescence. Cell Calcium, 10.1016/j.ceca.2017.04.001.

49. Streicher,F. and Jouvenet,N. (2019) Stimulation of Innate Immunity by Host and Viral RNAs. Trends Immunol., 10.1016/j.it.2019.10.009.

50. Patel,D.J.J. (2016) A structural perspective on readout of epigenetic histone and DNA methylation marks. Cold Spring Harb. Perspect. Biol., 10.1101/cshperspect.a018754.

